# The tyrosine phosphatase LAR acts as a receptor of the nidogen-tetanus toxin complex

**DOI:** 10.1101/2023.02.03.526966

**Authors:** Sunaina Surana, David Villarroel-Campos, Chiara Panzi, Sergey S. Novoselov, Sandy Richter, Giuseppe Zanotti, Giampietro Schiavo

## Abstract

Tetanus toxin is one of the most potent neurotoxins and is the causative agent of tetanus. This neurotoxin binds to the neuromuscular junction and, after internalisation into motor neurons, undergoes long-distance axonal transport and transcytosis into spinal cord inhibitory interneurons. Inside the cytoplasm of interneurons, the catalytic chain of the toxin blocks neurotransmitter release, leading to spastic paralysis. Whilst the effects of tetanus toxin intoxication have been extensively studied, the molecular composition of its receptor complex is still poorly understood. We have previously shown that the extracellular matrix proteins nidogens are essential for binding of the toxin to the neuromuscular junction. In this study, we show that the tyrosine phosphatase LAR interacts with the nidogen-tetanus toxin complex and enables its uptake into motor neurons. Binding of LAR to the toxin complex is mediated by its fibronectin III domains, which we have harnessed to inhibit tetanus toxin entry into motor neurons. Surprisingly, this function of LAR is independent of its role in regulating the neurotrophic activity of the TrkB receptor, which has previously been shown to augment the axonal transport of signalling endosomes containing tetanus neurotoxin. These findings identify a multi-subunit complex acting as a protein receptor for tetanus neurotoxin, and demonstrate a novel endocytic trafficking route for extracellular matrix proteins in neurons. Our study paves the way for dissecting the molecular mechanisms that control the recognition and uptake of physiological ligands and pathological proteins at the neuronal plasma membrane, as well as their targeting to the axonal retrograde pathway for long-distance transport within the nervous system.

## Introduction

Tetanus neurotoxin (TeNT) is one of the most toxic molecules identified to date. Produced by the anaerobic, Gram-positive bacterium *Clostridium tetani*, TeNT causes tetanus, a neuroparalytic syndrome characterised by lockjaw, opisthotonus, muscle stiffness and increasingly painful spasms, ultimately leading to respiratory failure and death (Farrar *et al*, 2000). Despite the availability of an effective vaccine and various antitoxin preparations, tetanus is a leading cause of mortality in neonates and unvaccinated adults in developing countries due to limited resources, lack of enforcement of appropriate public healthcare measures and scarce availability of treatments (Thwaites *et al*, 2015; Pirazzini *et al*, 2022).

TeNT is formed by three modular domains, each of which is essential for intoxication of the nervous system. In the active toxin, these domains are arranged in a heavy chain (H chain) and a catalytic light chain (L chain). Generated from a single polypeptide by proteolytic cleavage, both subunits remain associated *via* non-covalent interactions and a conserved inter-chain disulphide bond. The H chain is further subdivided into two regions: an amino terminal (H_N_) and a carboxy terminal (H_C_) domain, which are responsible for membrane translocation and receptor binding, respectively (Schiavo *et al*, 2000; Surana *et al*, 2018). After bacterial spore germination and TeNT production, the active toxin accumulates in the synaptic space at the neuromuscular junction (NMJ) and binds to the plasma membrane of motor neurons with sub-nanomolar affinity by virtue of its H_C_ domain (Pirazzini *et al*, 2016). This results in rapid internalisation of the neurotoxin, followed by its long-distance, retrograde transport towards the neuronal cell body in the spinal cord (Salinas *et al*, 2010). TeNT then undergoes trans-synaptic transfer into inhibitory interneurons, where the H_N_ domain drives the pH-dependent translocation of the L chain from the endocytic lumen of synaptic vesicles into the cytosol (Pirazzini *et al*, 2016). The L chain, which possesses zinc protease activity, cleaves the synaptic vesicle protein synaptobrevin, leading to a cessation of neurotransmitter release (Schiavo *et al*, 1992, 1994). This perturbs the balance of excitatory and inhibitory inputs to motor neurons, leading to motor neuron hyperactivity and spastic paralysis (Schiavo *et al*, 2000; Surana *et al*, 2018).

Given the high neuro-specificity and extreme toxicity of TeNT, several studies have endeavoured to identify the presynaptic receptors responsible for its entry into motor neuron terminals. The C-terminal region of the H_C_ domain (H_CC_) was found to bind with high affinity to polysialogangliosides of the G1b subgroup (Chen *et al*, 2009). This interaction, which takes place *via* two sialic acid-binding pockets, is essential for TeNT intoxication since site-directed mutagenesis of key residues within these sites led to a dramatic loss of binding to rat brain synaptosomes as well as abrogation of toxicity in a phrenic nerve-diaphragm preparation (Rummel *et al*, 2003). However, whereas polysialogangliosides are enriched in the neuronal plasma membrane, they are not exclusively present on the presynaptic surface of motor neurons. Additionally, membrane binding of TeNT was found to be protease-sensitive (Pierce *et al*, 1986; Lazarovici & Yavin, 1986). This, together with the finding that the H_C_-GT1b interaction displays a higher binding constant *in vitro* than that observed for H_C_ binding to rat brain synaptosomes (Rummel *et al*, 2003), suggested that instead of relying solely on polysialogangliosides, TeNT binding to motor neurons also requires a specific membrane receptor (Montecucco *et al*, 2004).

Early studies indicated that the protein receptor for TeNT was a glycophosphoinositol (GPI)-anchored protein resident in lipid microdomains, since treatment of mouse spinal cord cells with phosphatidylinositol-specific phospholipase C inhibited TeNT-induced cleavage of synaptobrevin (Munro *et al*, 2001). This was supported by the observation that gangliosides accumulate in lipid rafts (Simons & Toomre, 2000). These observations led to the identification of Thy-1 as a TeNT-binding protein in glycolipid-enriched membrane regions. However, independent evidence indicated that Thy-1 was not the neuronal receptor for TeNT since H_C_ binding and internalisation in Thy-1 knockout mice was indistinguishable from wild-type mice (Herreros *et al*, 2001). More recently, the extracellular matrix (ECM) proteins nidogens (also called entactins) were identified as co-receptors of TeNT at the NMJ. The H_C_ domain was found to bind preferentially to nidogen-rich regions of the NMJ, and both nidogen-1 and −2 were found to directly interact with TeNT. Furthermore, peptides derived from nidogen-1 were found to inhibit membrane binding and uptake of TeNT in motor neurons, which prevented the appearance of spastic paralysis in intoxicated mice (Bercsenyi *et al*, 2014). However, the precise identity of the membrane receptor that engages with the nidogen-TeNT complex and ferries it into motor neurons remains unknown.

Several lines of evidence suggest that the tyrosine phosphatase LAR (leukocyte common antigen-related protein) plays a role in the internalisation of TeNT. O’Grady and colleagues have shown that the nidogen-laminin complex interacts with recombinant LAR *in vitro*, a result supported by the observation that the *Caenorhabditis elegans* orthologue of LAR, *ptp-3*, was found to genetically interact with nidogen-1 (O’Grady *et al*, 1998; Ackley *et al*, 2005). Furthermore, LAR is sequestered in lipid microdomains on the plasma membrane and co-localises with caveolin-enriched fractions (Caselli *et al*, 2002). LAR has also been linked to trophic pathways in the nervous system, including in the regulation of BDNF-dependent signalling of its receptor TrkB in hippocampal neurons (Yang *et al*, 2006). Concomitantly, TeNT triggers TrkB phosphorylation, activates downstream Akt and ERK signalling (Gil *et al*, 2003; Calvo *et al*, 2012) and shares retrograde signalling endosomes with BDNF and its receptors TrkB and p75^NTR^ during its axonal journey to the spinal cord (Lalli & Schiavo, 2002; Deinhardt *et al*, 2006). Altogether, this evidence highlights functional links between the nidogen-TeNT complex and LAR.

LAR, along with PTPRδ and PTPRσ, belongs to the type IIa family of transmembrane receptor-type protein tyrosine phosphatases (RPTPs). These proteins contain three extracellular immunoglobulin-like (Ig) domains, followed by four to eight fibronectin III (FNIII) domains, depending on alternative splicing. The intracellular subunit contains two phosphatase domains: D1 and D2, of which only the D1 domain is catalytically active (**Fig 1A**). Mature RPTPs undergo constitutive proteolysis between the FNIII and D1 domains, generating an extracellular subunit that is non-covalently bound to the phosphatase domaincontaining moiety (Takahashi & Craig, 2013). LAR has been shown to regulate cellular adhesion and signalling, thus playing important roles in the nervous system, including synapse formation and stabilisation, axon guidance as well as sprouting and innervation in cholinergic neurons (Cornejo *et al*, 2021).

**Figure 1.**
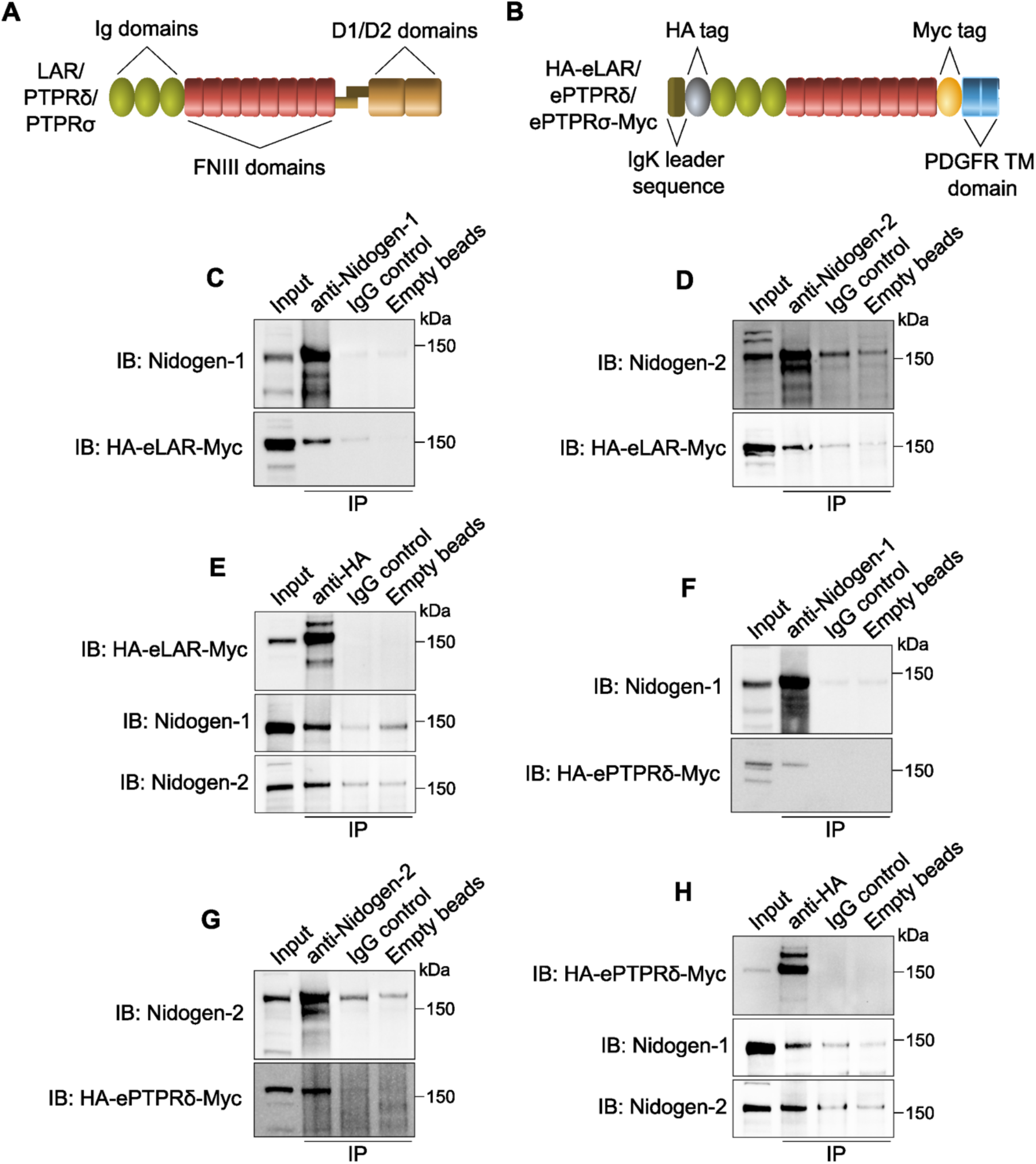
Nidogens bind to the receptor-type protein tyrosine phosphatases (RPTPs) LAR and PTPRδ. **(A)** Schematic showing the domain organisation of the LAR family of RPTPs. Full-length LAR, PTPRδ and PTPRσ contain three extracellular immunoglobulin-like (Ig) and eight fibronectin III (FNIII) domains, as well as two intracellular protein tyrosine phosphatase (D1 and D2) domains. RPTPs undergo proteolytic processing between the FNIII and phosphatase domains; this generates an extracellular subunit that remains non-covalently bound to the intracellular phosphatase subunit. **(B)** Schematic diagram of human LAR, PTPRδ and PTPRσ chimeric proteins used for co-immunoprecipitations. The extracellular domain of each of these RPTPs was fused to the murine Igκ-chain leader sequence and haemagglutinin (HA) tag at the N-terminus; the C-terminus, on the other hand, was fused to the platelet-derived growth factor receptor transmembrane (PDGFR TM) domain and a Myc tag. **(C-E)** Nidogen-1 and −2 directly interact with the extracellular domain of LAR in the presence of VSVG-H_C_T. Nidogen-1 (C) and nidogen-2 (D) were immunoprecipitated from N2a cell lysates, and co-immunoprecipitates were probed using an anti-HA antibody. Conversely, HA-eLAR-Myc was immunoprecipitated using an anti-HA antibody, followed by the detection of nidogens (E). Non-specific antibodies bound to beads and empty beads were used as controls; 5% input was loaded. **(F-H)** In the presence of H_C_T, PTPRδ is a binding partner of both nidogen-1 and −2. Nidogen-1 (F), nidogen-2 (G) and HA-ePTPRδ-Myc (H) were immunoprecipitated from lysates of N2a cells, following which coimmunoprecipitates were probed using an appropriate antibody (anti-HA for nidogen immunoprecipitations, and anti-nidogen for HA-eLAR-Myc immunoprecipitations). Non-specific antibodies bound to beads and empty beads were used as negative controls; 5% input was loaded.

In this study, we report that LAR is a component of the nidogen-TeNT complex and enables its internalisation in motor neurons. We show that depletion of LAR is sufficient to abrogate neuronal uptake of the H_C_ domain of TeNT (henceforth referred to as H_C_T). Binding of LAR to the nidogen-TeNT complex is mediated by specific fibronectin III domains, which we leveraged as tools to inhibit H_C_T entry into motor neurons. Importantly, this function of LAR is independent of its role in regulating the neurotrophic activity of TrkB. Taken together, our results define the physiological receptor for TeNT on the neuronal plasma membrane and yield important insights into the mechanisms controlling the recognition and internalisation of physiological ligands and toxins, such as TeNT, into the nervous system.

## Results

### Nidogens interact with the receptor-type protein tyrosine phosphatases LAR and PTPRδ

The starting point of our investigation was the observation that TeNT is critically dependent on nidogens for its binding to the mammalian NMJ, together with the previously reported *in vitro* interaction between LAR phosphatase and the nidogen-laminin complex (O’Grady *et al*, 1998; Bercsenyi *et al*, 2014). For LAR to act as the TeNT receptor at the NMJ, thus enabling its internalisation into motor neurons, the extracellular domain of LAR should be able to bind the TeNT-nidogen complex. To test this hypothesis, we performed co-immunoprecipitation experiments between the extracellular subunit of LAR and nidogens in the presence of H_C_T. The Ig and FNIII domains of human LAR (eLAR) were fused to a haemagglutinin (HA) tag at the N-terminus; the C-terminus was fused to a Myc tag and the transmembrane domain of the platelet-derived growth factor (PDGF) receptor (**Fig 1B**). When this fusion protein (HA-eLAR-Myc) was transiently expressed in mouse neuroblastoma-2a (N2a) cells, we found that it was retained on the membrane by virtue of its transmembrane domain (**Supplementary Fig 1A)**. Similarly, when mouse nidogen-1 or −2 were expressed in N2a cells, they were secreted into the extracellular medium and then bound to the cell surface (**Supplementary Fig 1A**). When recombinant nidogen-1 was co-expressed with HA-eLAR-Myc in N2a cells and immunoprecipitated in the presence of H_C_T, we found that eLAR was specifically associated with the beads containing anti-nidogen-1 antibodies (**Fig 1C**). Similarly, when nidogen-2 was co-expressed with HA-eLAR-Myc and subjected to immunoprecipitation, it robustly bound to the extracellular domain of LAR, as shown by the comparison with beads coated with non-immune mouse IgG or empty bead controls (**Fig 1D**). The direct interaction between nidogens and LAR was further confirmed when reverse co-immunoprecipitations were performed. Indeed, when HA-eLAR-Myc was immunoprecipitated using an anti-HA antibody, it co-precipitated both nidogen-1 and −2 (**Fig 1E**). This fits well with our unpublished results obtained *via* a proximity biotinylation approach in which LAR was identified as a potential interacting partner of H_C_T in signalling endosomes. In this approach, H_C_T was fused to a promiscuous biotin ligase (Roux *et al*, 2012), and allowed to be taken up in signalling endosomes upon incubation with embryonic stem cell-derived motor neurons. This led to enzyme-catalysed biotinylation, and subsequent identification by mass spectrometry, of proteins present within a 10-15 nm radius of the H_C_T fusion protein in neuronal endosomes, one of which was LAR (SS Novoselov and G Schiavo, unpublished data). Taken together, these data suggest that there is a direct physical interaction between nidogens and LAR, which is preserved upon their entry into signalling endosomes.

Since LAR, PTPRδ and PTPRσ share a high degree of homology and structural similarity (**Fig 1A**), we tested whether PTPRδ and PTPRσ also bind to nidogens. As described above, we co-expressed HA-ePTPRδ-Myc and nidogens in N2a cells and found that the extracellular domain of PTPRδ coimmunoprecipitated with both nidogen-1 and −2 in cell extracts (**Fig 1F, G; Supplementary Fig 1A**). Similarly, when HA-ePTPRδ-Myc was immunoprecipitated in the presence of H_C_T, it was found to contain both nidogens as interacting partners (**Fig 1H**). In contrast, recombinant HA-ePTPRσ-Myc could not be co-immunoprecipitated with either nidogen-1 or −2, suggesting a lack of interaction between nidogens and PTPRσ (**Supplementary Fig 1B, C**).

### LAR is co-distributed with the nidogen-H_C_T complex in signalling endosomes of motor neurons

After confirming the direct association between LAR and nidogens, we wanted to assess whether the nidogen-H_C_T complex and LAR are internalised and transported together in signalling endosomes. Primary motor neurons were incubated with AlexaFluor 647-labelled H_C_T (H_C_T-647) and an antibody against nidogen-2, which were allowed to internalise at 37°C. After removing the surface-bound probes using a mild acidic wash, the total cellular pool of LAR was detected using an anti-LAR antibody (Bercsenyi *et al*, 2014). After immunostaining with appropriate fluorescent secondary antibodies, we observed that endogenous LAR is present in the cell body as well as axons of mouse motor neurons. Interestingly, the axonal population of

LAR displays a punctate pattern reminiscent of endosomal compartments. Crucially, these LAR puncta co-localise with H_C_T and nidogen-2 in the axonal network and cell bodies, with many of them being triple positive (**Fig 2A, B**). This result strongly suggests that all three proteins were co-internalised and underwent long-distance retrograde transport in signalling endosomes. These observations are in line with our previous study that reported the presence of LAR in H_C_T-containing signalling endosomes purified from mouse embryonic stem cell-derived motor neurons (Debaisieux *et al*, 2016). Upon analysing fluorescence intensities of nidogen-2 and LAR in individual neurites in these cultures, we found that cells with higher levels of LAR contained, on average, higher levels of nidogen-2 in signalling endosomes, as shown by the significant correlation between the levels of internalised nidogens and endogenous LAR (Spearman coefficient = 0.497; **Fig 2C**). This correlation was stronger for internalised H_C_T and LAR, with a correlation coefficient of 0.743, indicating that cellular LAR levels directly correspond to endosomal levels of the nidogen-H_C_T complex (**Fig 2C**).

**Figure 2.**
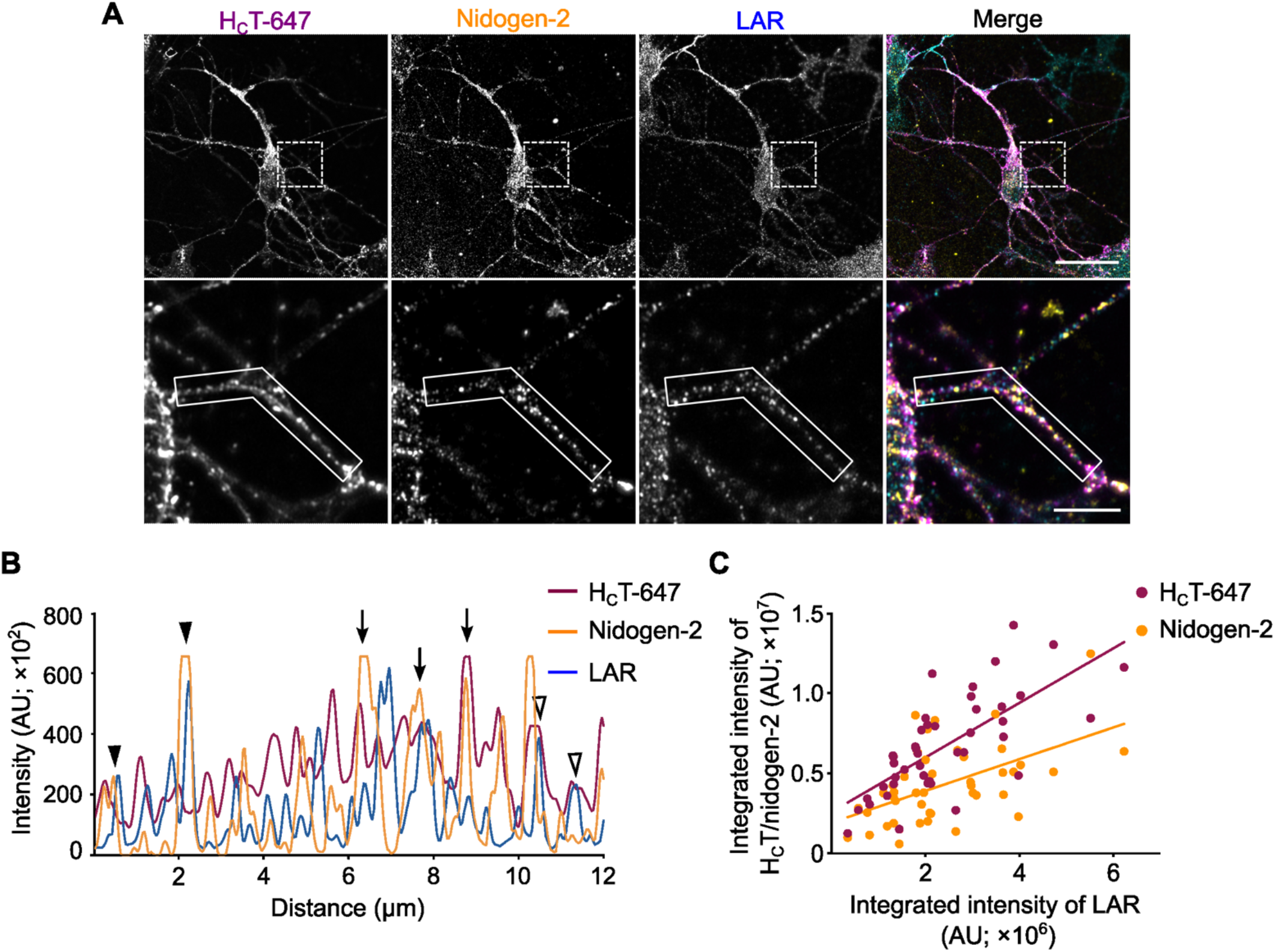
Endogenous LAR co-localises with the nidogen-H_C_T complex in signalling endosomes. **(A)** Representative immunofluorescence images of mouse motor neurons treated with H_C_T-647 and labelled with antibodies against internalised nidogen-2 and total LAR. Images have been pseudo-coloured in magenta (H_C_T-647), yellow (nidogen-2) and cyan (LAR). Selected region in the upper panel has been magnified in the lower panel. Scale bar: 20 μm (top panel) and 5 μm (bottom panel). **(B)** Graph showing overlapping intensity profiles of H_C_T-647, nidogen-2 and LAR in an axonal segment (boxed region in the lower panel of (A). Empty arrowheads point to colocalised H_C_T and LAR organelles, arrowheads denote co-localised nidogen-2 and LAR puncta, while filled arrows represent puncta containing H_C_T, nidogen-2 and LAR. **(C)** Fluorescence intensities of both H_C_T-647 and nidogen-2 exhibit significant correlation with LAR fluorescence intensities in motor neurons (n = 46 neurites; Spearman coefficient 0.743 and 0.497, and *P* < 0.0001 and 0.0004, for LAR-H_C_T and LAR-nidogen-2, respectively).

### Depletion of LAR inhibits H_C_T entry in motor neurons

If the nidogen-TeNT complex were indeed a ligand for the surface LAR receptor, we reasoned that a decrease in LAR levels would lead to an inhibition of H_C_T binding and uptake. To assess this, we transduced ventral horn cultures with lentiviruses encoding short hairpin RNAs (shRNAs) against mouse LAR. After 48 h of viral transduction, we found that two independent shRNAs were able to reduce endogenous LAR levels by ~70%, compared to empty vector and scrambled controls (**Fig 3A, B**). Critically, in this time window, we did not observe any overt alterations in neuronal survival or gross morphology (**Fig 3C**). We confirmed the specificity of LAR knockdown by assessing levels of PTPRδ and PTPRσ in these cultures and found that unlike LAR, the levels of these proteins remained unchanged after LAR knockdown (**Fig 3A; Supplementary Fig 2A, B**). When these cultures were treated with H_C_T-647 for 1 h at 37°C, we found that transduced motor neurons, which were identified by the expression of green fluorescent protein (GFP), exhibited a significant decrease in H_C_T internalisation (~40%), compared to controls (**Fig 3C, D**). The decrease in H_C_T uptake was observed using both shRNAs, suggesting that this effect is specific and due to LAR downregulation rather than potential off-target effects. To confirm this conclusion, motor neurons transduced with LAR shRNA#2 were magnetofected with an HA-eLAR-Myc-expressing plasmid (**Fig 1B**). After 24 h, cultures were incubated with H_C_T-647 for 1 h, fixed, immunostained for GFP, HA and βIII tubulin, and imaged. We found that neurons expressing both shRNA#2 and recombinant eLAR, which were identified by the presence of GFP and HA, respectively, showed a significant increase in endocytosis of H_C_T, compared to cultures expressing shRNA#2 alone, and was commensurate with H_C_T levels observed in cultures treated with scrambled shRNAs (**Fig 3C, D**). This rescue using the extracellular LAR domain confirms that the decrease in H_C_T endocytosis observed upon LAR knockdown is specific and that levels of H_C_T internalisation are directly correlated with neuronal LAR levels.

**Figure 3.**
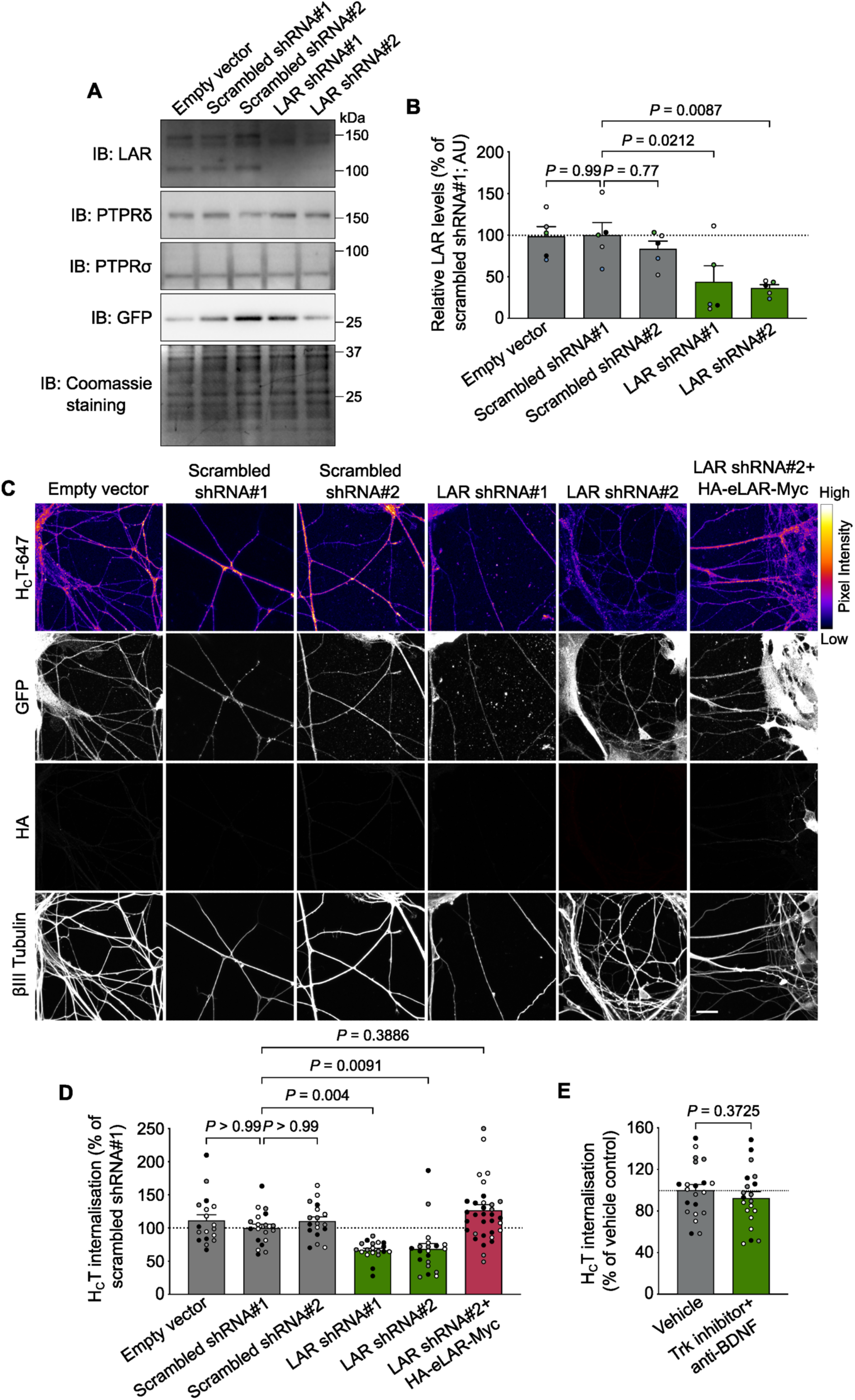
Depletion of LAR causes a decrease in H_C_T internalisation in motor neurons. **(A)** Representative western blots for estimating the levels of LAR, PTPRδ and PTPRσ in lysates of ventral horn cultures transduced with lentiviruses encoding short hairpin RNAs (shRNAs) against murine LAR. Lentiviruses carrying an empty vector and two scrambled shRNAs were used as negative controls, whereas GFP was used as a transduction reporter. Coomassie R-250 staining was used to estimate protein loading. **(B)** LAR quantification shown in (A). Data are presented as a percentage of LAR levels in ventral horn cultures treated with scrambled shRNA#1 (n = 5 independent experiments; error bars indicate s.e.m.). Results were tested for statistical significance using one-way analysis of variance (ANOVA; *P* = 0.004), followed by Dunnett’s *post-hoc* test. **(C)** Representative immunofluorescence images showing internalised H_C_T-647 in motor neurons, following lentiviral-mediated knockdown of endogenous LAR, as well as its rescue by overexpression of recombinant HA-eLAR-Myc. Images in the H_C_T-647 panel have been colour mapped based on their intensities. GFP was used as a reporter of lentiviral transduction, while the HA tag was used to confirm expression of HA-eLAR-Myc. Lentiviruses carrying an empty vector and two scrambled shRNAs were used as negative controls. Scale bar: 20 μm. **(D)** Quantification of endocytosed H_C_T-647 shown in (C). Data are presented as a percentage of internalised H_C_T in neurons treated with scrambled shRNA#1 (n = 3 independent experiments; error bars indicate s.e.m.). Results were analysed for statistical significance using Kruskal-Wallis test (*P* < 0.0001), followed by Dunn’s multiple comparison test. **(E)** Levels of internalised H_C_T in motor neurons treated with the pan-Trk inhibitor PF-06273340 and an anti-BDNF antibody, compared to vehicle control (DMSO). Data are presented as a percentage of internalised H_C_T in neurons treated with DMSO alone (n = 3 independent experiments; error bars indicate s.e.m.). Results were tested for statistical significance using an unpaired *t*-test.

Tyrosine phosphatases such as LAR are known to regulate the phosphorylation of several tyrosine kinases, ultimately influencing their downstream signalling (Kulas *et al*, 1996; Sarhan *et al*, 2016). Accordingly, LAR has been previously shown to interact with the tyrosine kinase TrkB in hippocampal neurons, thus modulating its neurotrophic activity. Neurons devoid of LAR displayed a reduction in the BDNF-induced phosphorylation of TrkB, which led to a decrease in the phosphorylation of its downstream effectors Shc, Akt and ERK, ultimately translating into an increase in neuronal cell death (Yang *et al*, 2006). Based on this evidence, it was possible that the decrease in H_C_T uptake observed upon LAR knockdown in motor neurons was indirect and caused by a reduction in TrkB phosphorylation. To rule out this possibility, we tested the effect of PF-06273340, a highly potent and selective inhibitor of Trk receptors (Skerratt *et al*, 2016). Ventral horn cultures were treated with 100 nM of PF-06273340, together with an anti-BDNF antibody, for 30 min to completely abrogate TrkB signalling. HA-H_C_T was then added for 1 h, after which the cells were fixed, immunostained for βIII tubulin as well as the HA tag, and imaged.

We found that Trk inhibition had no overt effect on H_C_T internalisation under these experimental conditions, thus ruling out that the inhibition of nidogen-TeNT uptake observed upon LAR downregulation in motor neurons was mediated by an indirect effect on TrkB signalling (**Fig 3E; Supplementary Fig 2C**).

These results show that the internalisation of H_C_T requires the expression of LAR in motor neurons and that this role of LAR is independent of its modulatory function on TrkB. Taken together, these findings suggest that LAR acts as the cellular receptor for the nidogen-H_C_T complex.

### LAR-nidogen binding is mediated by the fibronectin III domains of LAR

Next, we wanted to characterize the interaction between LAR and nidogen at the molecular level using a protein truncation and co-immunoprecipitation approach. First, the extracellular domain of LAR was truncated into three fragments containing: i) the Ig domains (HA-LAR Ig1-3-Myc), ii) the first four FNIII domains (HA-LAR FNIII1-4-Myc) and, iii) FNIII domains five to eight (HA-LAR FNIII5-8-Myc) (**Fig 4A**). Similar to the HA-eLAR-Myc fusion protein, constructs were fused to the transmembrane domain of the PDGF receptor to enable their surface localisation. When these fusion proteins were expressed together with nidogen-2 in N2a cells and immunoprecipitated using anti-HA beads, we found that nidogen-2 could only be co-immunoprecipitated with constructs containing FNIII domains (**Fig 4B**). This result indicated that the LAR-nidogen interaction is mediated exclusively by the FNIII domains of LAR.

**Figure 4.**
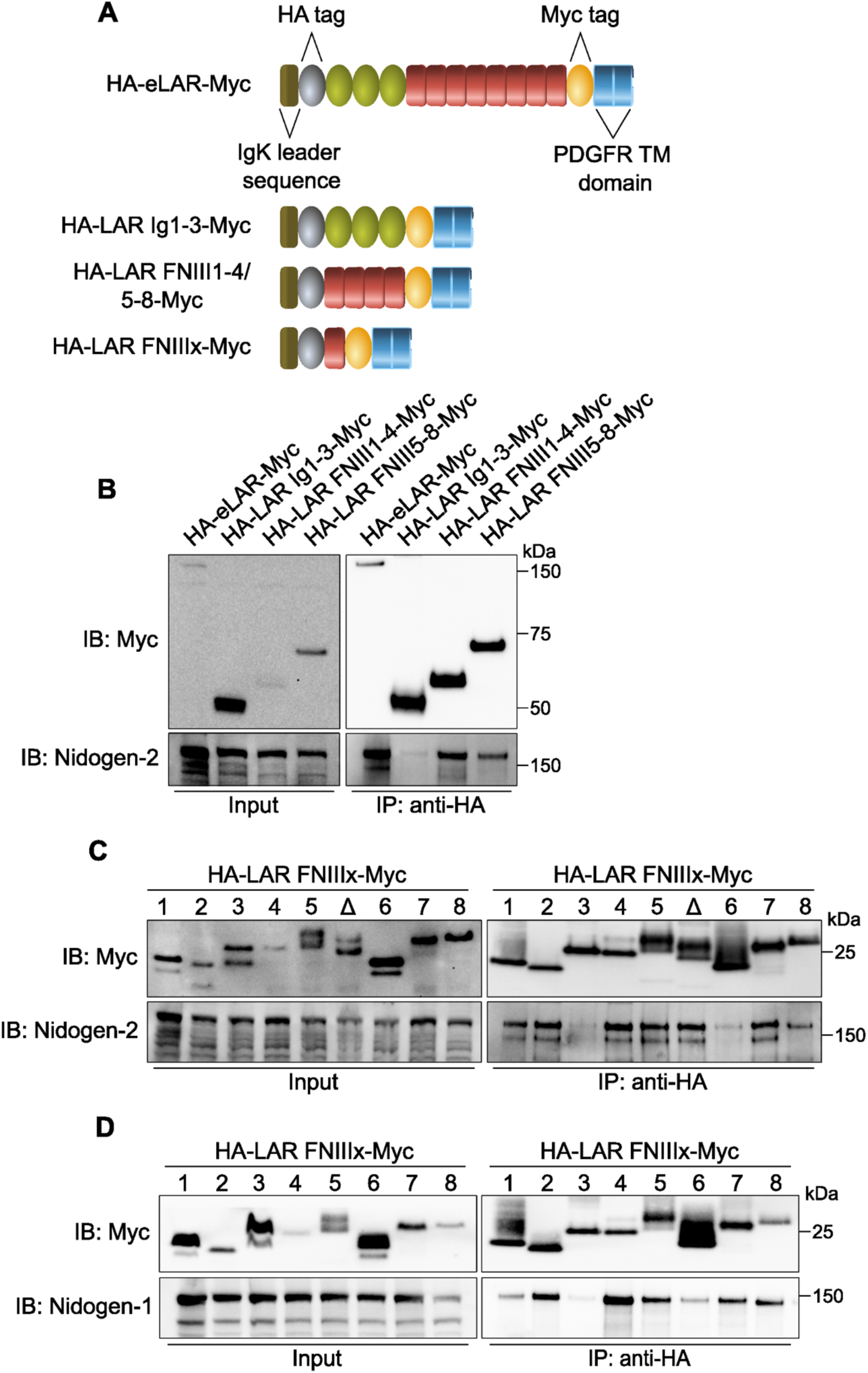
Nidogens bind to specific fibronectin III domains of LAR. **(A)** Schematics of LAR fragments used to identify the interacting domains between LAR and nidogens. Truncated proteins were fused to the murine Igκ-chain leader sequence and an HA tag at the N-terminus; the C-terminus was fused to the PDGFR transmembrane domain and a Myc tag. **(B)** Co-immunoprecipitation and western blot analysis of the interaction between nidogen-2 and HA-LAR Ig1-3-Myc, HA-LAR FNIII1-4-Myc and HA-LAR FNIII5-8-Myc, in the presence of VSVG-H_C_T. Immunoprecipitation was performed using an anti-HA antibody, and coimmunoprecipitated samples were probed using an anti-nidogen-2 antibody. The HA-eLAR-Myc fusion protein was used as a positive control and 5% input was loaded. **(C, D)** Western blots showing the interaction between individual LAR FNIII domains and nidogen-2 (C) or nidogen-1 (D), in the presence of VSVG-H_C_T. The Δ lane refers to co-immunoprecipitations performed using the 5^th^ FNIII domain without the MeC mini-exon. All immunoprecipitations were performed using an anti-HA antibody, whilst co-immunoprecipitates were probed using an appropriate anti-nidogen antibody. 5% input was loaded.

We then cloned all eight FNIII domains (HA-LAR FNIIIx-Myc) individually to identify the specific site(s) of interaction with nidogen-2 (**Fig 4A**). Co-expression and co-immunoprecipitation analyses demonstrated that the 2^nd^, 4^th^, 5^th^ and 7^th^ FNIII domains efficiently pull down nidogen-2 (**Fig 4C**). While the 1^st^ and 8^th^ FNIII domains could also co-precipitate nidogen-2, the immunoprecipitation yield was noticeably lower, suggesting that the interaction is likely to be weaker. In contrast, the 3^rd^ and 6^th^ FNIII domains only displayed non-specific binding of nidogen-2 to anti-HA beads (**Fig 4C**). Collectively, these results indicate that the association between LAR and nidogen is likely to be multivalent, *i.e*., mediated by interactions with multiple FNIII domains across the LAR extracellular domain.

LAR undergoes alternative splicing to generate cell type- and developmental stage-specific isoforms, which display unique interaction profiles. One of these variants, which is specifically expressed in the nervous system, is generated by the alternative splicing of a nine amino acid cassette in the 5^th^ FNIII domain (O’Grady *et al*, 1994; Zhang & Longo, 1995). The Saito group has previously shown that splicing of this mini-exon, termed MeC, is essential for the *in vitro* interaction between the 5^th^ FNIII domain and the laminin-nidogen complex (O’Grady *et al*, 1998). In light of this observation, we wanted to test whether the MeC cassette plays a similar role in the binding of LAR to the nidogen-H_C_T complex. We deleted this miniexon from the HA-LAR FNIII5-Myc construct and then assessed the ability of this variant to associate with nidogen-2. We found that nidogen-2 binds both splice variants of the 5^th^ FNIII domain, indicating that inclusion of this mini-exon does not alter the ability of LAR to interact with nidogen-2 (**Fig 4C**). This observation suggests that the mode of binding of LAR to the nidogen-TeNT complex is different from its interaction with the laminin-nidogen complex and is independent of the MeC mini-exon.

Finally, we wanted to test whether nidogen-1 mirrors the LAR-binding properties of nidogen-2. We performed co-immunoprecipitations with individual LAR FNIII domains and nidogen-1 under the same experimental conditions described for nidogen-2, and found that while the overall pattern of binding remained unchanged, there were subtle differences in the strength of the interactions. While the 2^nd^, 4^th^, 5^th^ and 7^th^ FNIII domains were the strongest interactors, the 1^st^ FNIII domain displayed a reduced ability to pull down nidogen-1 in the immunoprecipitate (**Fig 4D**). In contrast, the 8^th^ FNIII domain showed an increased affinity for nidogen-1 than nidogen-2 (**Fig 4D**). Akin to nidogen-2, nidogen-1 did not interact with the 3^rd^ and 6^th^ FNIII domains of LAR.

### Recombinant LAR fibronectin III domains halt the internalisation of the nidogen-H_C_T complex

Having mapped the interacting domains between LAR and nidogens, we sought to identify the shortest peptide sequences necessary for this interaction, with the aim of using these binding motifs to design competitive inhibitors disrupting the LAR-nidogen interaction. Using the multiple alignment software PRALINE, we aligned the sequences of the 2^nd^, 4^th^, 5^th^ and 7^th^ FNIII domains of human LAR and screened for amino acid conservation, as well as for similarities in hydrophobicity/hydrophilicity (Simossis & Heringa, 2005). Despite possessing a conserved secondary structure, none of these domains were found to contain conserved regions mediating binding to nidogens (**Supplementary Fig 3**).

In the absence of clear sequence similarities, together with the well documented modularity and functionality of individual fibronectin III domains, we decided to use full-length nidogen-binding domains of LAR as competitive inhibitors of the LAR-nidogen interaction (Petersen *et al*, 1983; Vilstrup *et al*, 2020).

To achieve this, we first cloned the 2^nd^, 4^th^, 5^th^ and 7^th^ FNIII domains of LAR into a bacterial expression vector, such that each recombinant protein is fused to a 6×His tag at the N-terminus and a FLAG tag at the C-terminus (**Supplementary Fig 4A**). Upon bacterial expression, each domain was purified using Ni^2+^-based affinity purification (**Supplementary Fig 4B-E; Supplementary Fig 5A-D**) (Vilstrup *et al*, 2020). We then used enzyme-linked immunosorbent assay (ELISA) to characterise the binding of each of these domains to full-length nidogen-2. Varying concentrations of each FNIII domain were added to 0.5 picomoles of recombinant mouse nidogen-2 in solution, which was then captured using an anti-nidogen-2 antibody. Subsequent complex detection showed that purified LAR FNIII domains bound to nidogen-2, indicating that they are indeed functional. Complex formation increased as a function of the *in vitro* FNIII domain concentration and followed a sigmoidal curve (**Fig 5**). Using this approach, we estimated that the average binding affinity of LAR FNIII domains to nidogen-2 was ~2 μM. Of these, the 5^th^ FNIII domain was the strongest binder, with an apparent binding affinity of ~1.4 μM, while the 7^th^ FNII domain bound with an apparent affinity of ~2.5 μM. The 2^nd^ and 4^th^ FNIII domains each displayed an apparent association constant of ~2 μM and ~1.9 μM, respectively (**Fig 5**).

**Figure 5.**
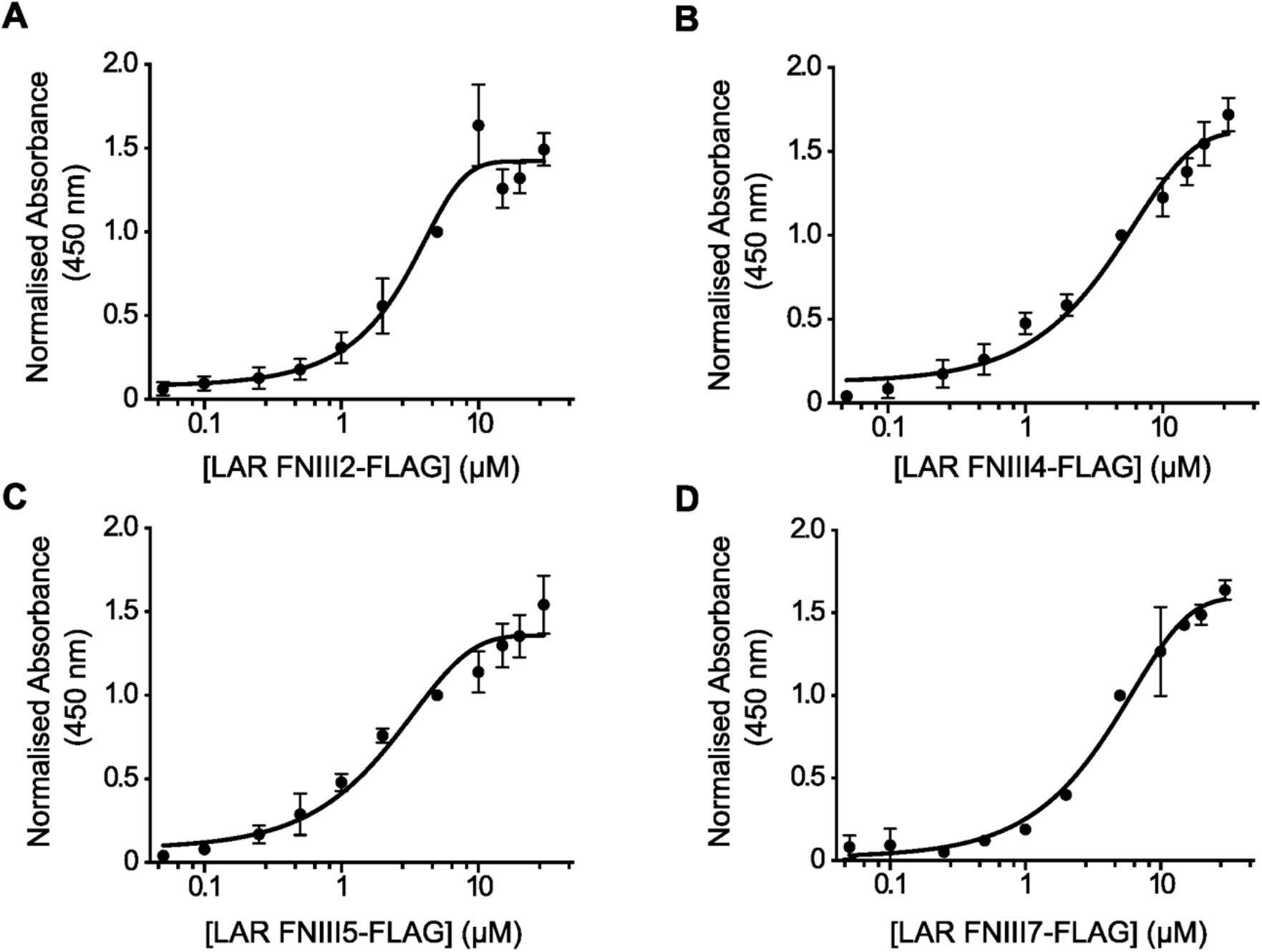
Dose-dependence of the interaction between nidogen-2 and soluble LAR FNIII2, FNIII4, FNIII5 and FNIII7 domains. **(A-D)** Plots showing the *in vitro* binding between nidogen-2 and purified LAR FNIII2-FLAG (A), LAR FNIII4-FLAG (B), LAR FNIII5-FLAG (C), and LAR FNIII7-FLAG (D). Serial dilutions of each LAR FNIII domain (50 nM-30 μM) were added to a fixed amount of immobilised nidogen-2 (0.5 picomoles), followed by addition of an anti-FLAG antibody to reveal complex formation using absorption spectroscopy. All datapoints were normalised to the absorbance obtained at the 5 μM LAR FNIII concentration (n = 3 independent experiments; error bars indicate s.e.m.).

We next tested whether these domains were able to individually bind to endogenous nidogens, potentially blocking the LAR-nidogen interaction and thus acting as competitive inhibitors of the uptake of the nidogen-H_C_T complex. Since our previous experiments had demonstrated an apparent binding affinity of ~2 μM, we decided to incubate motor neurons with H_C_T-555 and 20 μM of each purified FNIII domain for 30 min at 4°C. After media removal, motor neurons were transferred to 37°C for 45 min, which allows membranebound H_C_T to be internalised and undergo long-distance transport.

Firstly, we found that addition of FNIII recombinant domains did not affect cell health and morphology, as evidenced by βIII tubulin staining (**Fig 6A**). Addition of the 2^nd^, 4^th^ and 7^th^ FNIII domains led to a ~25% decrease in the uptake of nidogen-2 in axons, compared to buffer-treated controls (**Fig 6A, B**). However, the 5^th^ FNIII domain, which had the highest binding affinity to nidogen-2 among these domains, did not block nidogen-2 internalisation (**Fig 6B**). Importantly, these results were recapitulated upon quantification of H_C_T-555 uptake in these cells. Similar to nidogen-2, the 2^nd^, 4^th^ and 7^th^ FNIII domains led to a ~30% decrease in the uptake of H_C_T, while the 5^th^ FNIII domain did not show any effect (**Fig 6A, C**). These results suggest that whereas individual, recombinant LAR FNIII domains are capable of interacting with nidogens, their efficiency in interfering with the LAR-nidogen interaction in a cellular context is limited.

**Figure 6.**
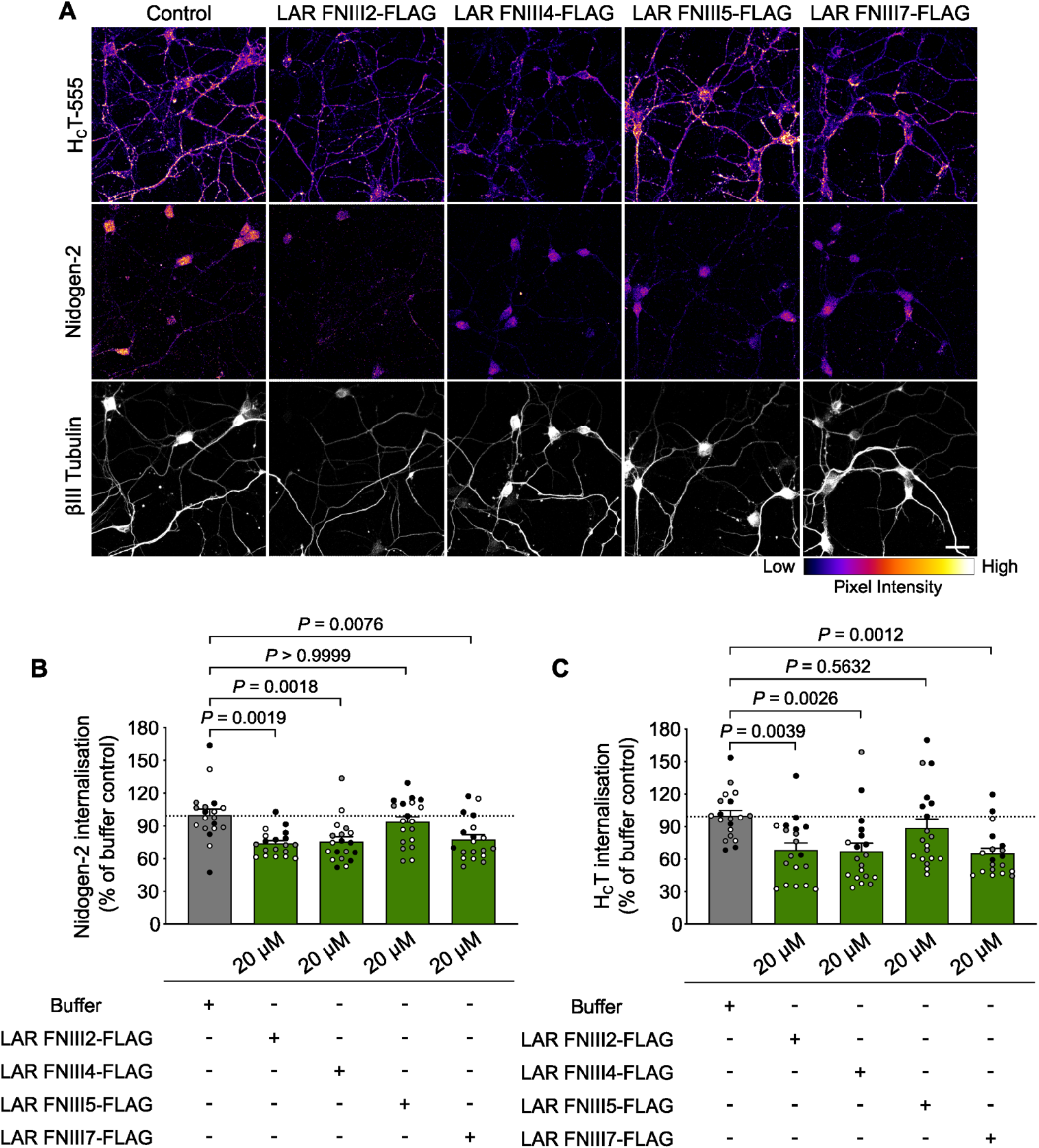
Recombinant LAR fibronectin III domains display a limited effect in inhibiting the binding of the nidogen-H_C_T complex to endogenous LAR. **(A)** Representative immunofluorescence images of motor neurons upon internalisation of H_C_T-555 and nidogen-2 in the presence of 20 μM recombinant, soluble LAR FNIII2, FNIII4, FNIII5 or FNIII7 domains. Images in the top two panels have been colour mapped based on their intensities. Scale bar: 20 μm. **(B, C)** Graph showing quantification of endocytosed nidogen-2 (B) and H_C_T-555 (C) shown in panel (A). Control refers to cultures treated with buffer alone. Data are presented as a percentage of internalised nidogen-2 or H_C_T in buffer-treated motor neurons (n = 3 independent experiments; error bars indicate s.e.m.). Results were tested for statistical significance using Kruskal-Wallis test (*P* = 0.0001), followed by Dunn’s *post-hoc* test (B) and one-way ANOVA (*P* = 0.0005), followed by Dunnett’s multiple comparisons test (C).

We next wanted to assess whether soluble FNIII domains added together can bind more efficiently to endogenous nidogens, thus abrogating the LAR-nidogen interaction. H_C_T-555 was added to ventral horn cultures in the presence of three concentrations of recombinant FNIII domains: 0.25 μM, 10 μM and 20 μM of each domain. After a 30 min pulse at 4°C, internalised H_C_T was chased for 45 min at 37°C, following which cells were fixed and immunostained for βIII tubulin and nidogen-2. We found that the amount of internalised nidogen-2 in cells incubated with 10 μM and 20 μM of recombinant FNIII domains was significantly lower than buffer-treated cells (**Fig 7A, B**). A decrease of ~40% was consistently observed when all four FNIII domains were added, and the extent of inhibition was higher than that observed using any single domain on its own. Interestingly, this decrease was observed even at the lower dose of each FNIII domain (0.25 μM), suggesting that the LAR-nidogen interaction might be governed by the combined avidity of individual FNIII domains.

**Figure 7.**
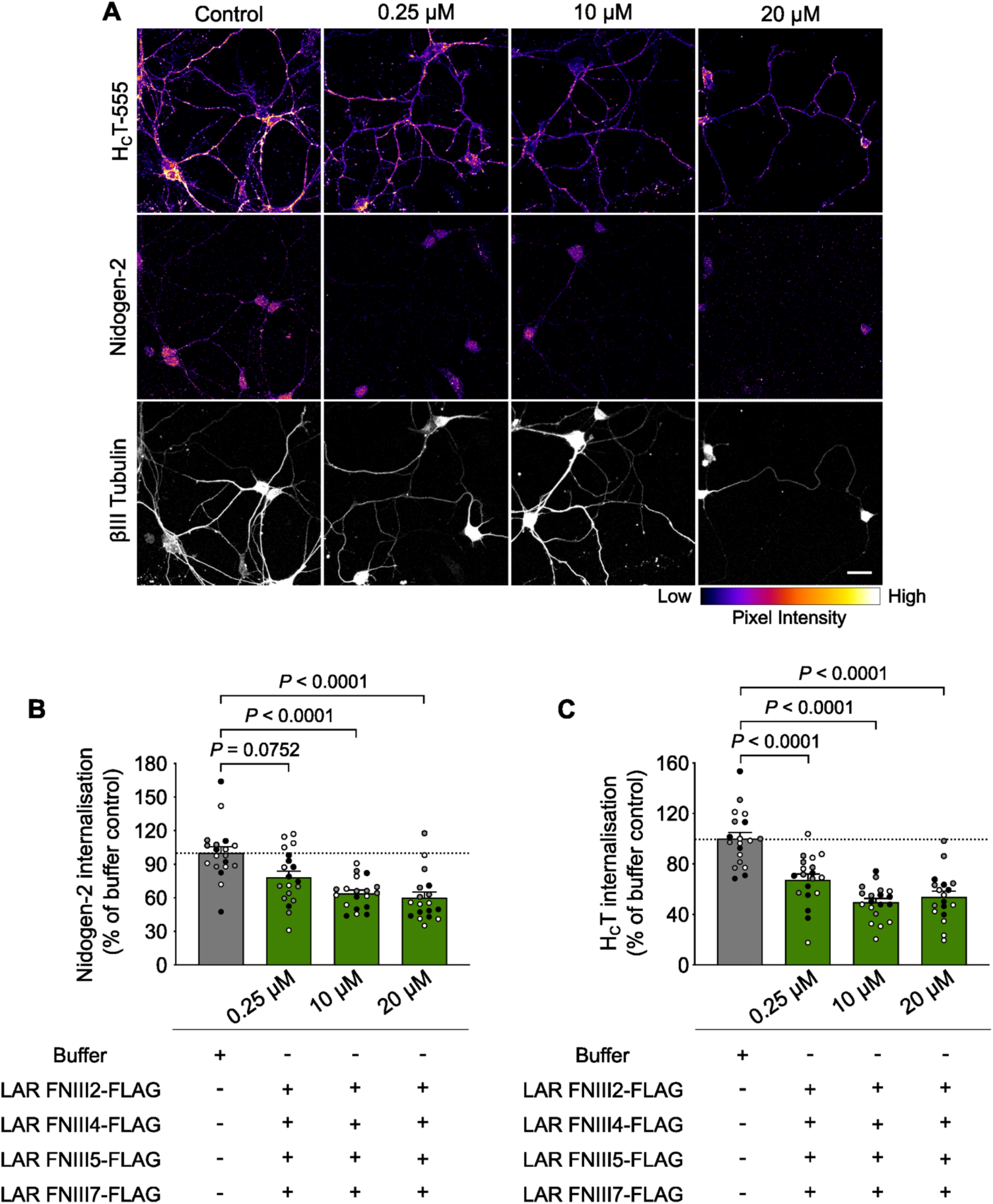
Multiple recombinant LAR fibronectin III domains block the nidogen-LAR interaction in motor neurons. **(A)** Representative immunofluorescence images of motor neurons upon internalisation of H_C_T-555 and nidogen-2 in the presence of 0.25 μM, 10 μM and 20 μM of each nidogen-binding FNIII domain (LAR FNIII2, FNIII4, FNIII5 and FNIII7). Images in the top two panels have been colour mapped based on their intensities. Scale bar: 20 μm. **(B, C)** Quantification of endocytosed nidogen-2 (B) and H_C_T-555 (C) shown in panel (A). Control refers to cultures treated with buffer alone. Data are presented as a percentage of internalised nidogen-2 or H_C_T in buffer-treated motor neurons (n = 3 independent experiments; error bars indicate s.e.m.). Results were analysed for statistical significance using Kruskal-Wallis test (*P* < 0.0001), followed by Dunn’s *post-hoc* test in (B) and one-way ANOVA (*P* < 0.0001), followed by Dunnett’s multiple comparisons test in (C).

When the intensity of internalised H_C_T was assessed in these cells, we found that its uptake was reduced by ~50% compared to controls (**Fig 7A, C**). The ability of soluble LAR FNIII domains to collectively abolish the binding of the nidogen-H_C_T complex to endogenous LAR and block its internalisation into motor neurons strongly indicates that LAR is the cellular receptor of the nidogen-TeNT complex in motor neurons.

## Discussion

TeNT is a highly potent toxin that causes spastic paralysis by inhibiting neurotransmission in spinal cord inhibitory interneurons. Entry into the central nervous system is achieved by targeting the mammalian NMJ, which leads to its internalisation into motor neurons and subsequent transcytosis into interneurons (Schiavo *et al*, 2000; Surana *et al*, 2018). Due to the high toxicity of TeNT and its causal role in human disease, a precise understanding of the physiological determinants enabling its entry into the nervous system is urgently needed. Previous studies have shown that binding of TeNT to the NMJ requires the presence of surface polysialogangliosides. However, protein(s) also play an essential role in concentrating TeNT at the NMJ and enabling its entry into motor neurons (Montecucco *et al*, 2004). Indeed, the ECM proteins nidogens were found to bind TeNT, thereby suggesting a mechanism involving the capture of TeNT at the NMJ and facilitating its neuronal entry (Bercsenyi *et al*, 2014). However, the identity of the membrane receptor that binds to the nidogen-TeNT complex on the surface of motor neurons and targets it to long-distance axonal transport is currently unknown. In this study, we show that the transmembrane tyrosine phosphatase LAR directly interacts with the nidogen-TeNT complex and ferries it into motor neurons, thus acting as its neuronal receptor.

LAR, and its homologues PTPRδ and PTPRσ, have been described as synaptic organisers that play important roles in the developing and mature nervous system, including axon guidance and neurite extension, as well as synapse formation, differentiation, and plasticity (Cornejo *et al*, 2021). The receptorlike extracellular domain has been reported to interact with a variety of *trans*-synaptic ligands and ECM molecules (such as netrin-G ligand-3, heparan sulphate proteoglycans and laminin, in the case of LAR), thereby modulating cell adhesion (O’Grady *et al*, 1998; Johnson & Van Vactor, 2003; Woo *et al*, 2009). On the other hand, the catalytic phosphatase subunit regulates the phosphorylation status and activity of many cellular proteins, including signalling molecules such as liprins (Kulas *et al*, 1996; Um & Ko, 2013; Sarhan *et al*, 2016). It is currently hypothesised that LAR-RPTPs act as molecular linkers that couple ECM components with downstream signalling cascades in the nervous system.

The starting point of our study was the previously reported interaction between LAR and nidogens. While Ackley and co-workers had shown a genetic interaction between these proteins, the Saito laboratory demonstrated that the laminin-nidogen complex acts as a ligand for LAR (O’Grady *et al*, 1998; Ackley *et al*, 2005). However, neither study showed a direct interaction between these molecules. Here, we have demonstrated that LAR binds to both nidogen-1 and −2 in the presence of H_C_T. These experiments, which were performed using only the extracellular portion of LAR, suggest that the intracellular phosphatase domain is dispensable for this interaction. This is in line with the observation that several RPTPs carry out their extracellular functions independently of their phosphatase domains (Young *et al*, 2021). It is, however, currently unclear whether nidogen binding modulates the phosphatase activity of LAR or changes the specificity of its downstream targets. In addition, we observed that whilst the LAR homologue PTPRδ also binds to nidogens, PTPRσ does not, confirming previous reports that these proteins are highly selective in their binding properties (Coles *et al*, 2015), in spite of their sequence and structural similarities.

Clostridial neurotoxins, such as TeNT, are known to hijack endogenous trafficking pathways to gain access to the nervous system and evade intracellular degradation (Surana *et al*, 2018). We have previously reported that, after binding to nidogen-rich regions at the NMJ, TeNT accomplishes its journey from the NMJ to the spinal cord by hitchhiking on signalling endosomes. These endocytic organelles contain several ligand-receptor complexes, including the neurotrophin BDNF and its receptors TrkB and p75^NTR^ (Lalli & Schiavo, 2002; Deinhardt *et al*, 2006). During its internalisation and intracellular transport, TeNT triggers Trk receptor phosphorylation, leading to initiation of downstream signalling cascades *via* activation of phospholipase Cγ-1 and phosphatidylinositol 3-kinase (Gil *et al*, 2003; Calvo *et al*, 2012). Concomitantly, multiple studies have reported the localisation of LAR-RPTPs at synaptic regions, with LAR particularly enriched at excitatory synapses and NMJs (Kaufmann *et al*, 2002; Ackley *et al*, 2005; Dunah *et al*, 2005). Interestingly, Ackley and colleagues have shown that synaptic accumulation of LAR is dependent on the presence of nidogens in the neuronal basement membrane (Ackley *et al*, 2005). LAR regulates vesicular trafficking by recruiting synaptic vesicles to active zones and coupling exo-endocytosis (Takahashi & Craig, 2013). All three LAR-RPTPs are present in H_C_T-containing signalling endosomes in motor neurons, with both LAR and PTPRδ regulating the BDNF-dependent phosphorylation and activation of TrkB (Yang *et al*, 2006; Debaisieux *et al*, 2016; Tomita *et al*, 2020). Furthermore, our experiments have shown that LAR presents a punctate pattern in motor neurons and is co-distributed with H_C_T and nidogen-2, indicating their presence in shared axonal carriers along common intracellular trafficking routes. In keeping with a putative role of LAR as a receptor for the nidogen-TeNT complex, LAR knockdown was sufficient to inhibit H_C_T internalisation in motor neurons. This result was confirmed with two independent shRNAs and rescued by expression of an shRNA-resistant extracellular subunit of LAR, strongly indicating that this effect was specific for LAR and not caused by off-target effects. We also observed that H_C_T and nidogen-2 uptake was commensurate with cellular LAR levels, further lending support to our hypothesis. This receptor function of LAR is independent of its role in regulating the phosphorylation status of the TrkB receptor, since incubation with a validated TrkB inhibitor had no effect on H_C_T uptake in neurons under our experimental conditions.

In an effort to better understand the LAR-nidogen interaction, we truncated the LAR extracellular domain and performed co-immunoprecipitations with full-length nidogens. We found that rather than binding to a single site, nidogens make multiple contacts along the length of the LAR extracellular domain. All the identified areas lie on FNIII domains, with the strongest interaction sites present on the 2^nd^, 4^th^, 5^th^ and 7^th^ FNIII domains, pointing to a multivalent interaction. However, we are unable to posit whether this binding is cooperative, that is, whether binding of one FNIII domain to a nidogen molecule facilitates binding of the remaining sites.

The stoichiometry of the LAR-nidogen interaction also remains unclear. Despite displaying the same structural fold, we were unable to discover any significant sequence similarity between the 2^nd^, 4^th^, 5^th^ and 7^th^ FNIII domains, barring amino acids that are necessary for fibronectin III domain folding. This is in stark contrast to the highly conserved RGD motif in the 10^th^ FNIII domain of the ECM protein fibronectin, which is essential for its interaction with integrins *in vivo* (Takahashi *et al*, 2007). Nonetheless, the relative *in vitro* binding affinities of recombinant nidogen-2 and LAR FNIII domains yielded an average value ~2 μM. This is consistent with the *in vitro* binding affinities of other surface protein-ECM pairs, which have been reported to rely on avidity of multivalent interactions rather than affinities of single binding events (Wright, 2009). Moreover, these experiments were performed using soluble LAR FNIII domains untethered to a membrane; this contrasts with the interaction taking place in cells where anchoring of FNIII domains on the plasma membrane would result in increased local concentrations of receptor molecules, thereby increasing the binding affinity. Interestingly, LAR is sequestered in caveolin-containing membrane microdomains, which are known to be enriched in cholesterol, glycosphingolipids and sphingomyelin (Caselli *et al*, 2002). Interactions with its extracellular binding partners triggers local clustering of LAR on the membrane, leading to formation of higher-order complexes (Um *et al*, 2014; Won *et al*, 2017; Xie *et al*, 2020). This molecular clustering is similarly observed for polysialogangliosides in lipid microdomains, which act as primary receptors as well as concentrating platforms for TeNT (Prinetti *et al*, 2000; Herreros *et al*, 2001). LAR is thus ideally positioned to bind to TeNT, along with polysialogangliosides and nidogens, and initiate its internalisation and delivery to the central nervous system.

The validity of our hypothesis was further confirmed by the efficient abrogation of H_C_T uptake by the addition of soluble LAR FNIII domains to motor neuron cultures. If LAR is indeed a surface receptor for the nidogen-TeNT complex, we reasoned that soluble FNIII domains, when added in excess, would outcompete the endogenous LAR-nidogen interaction and thus disrupt it, leading to a decrease in endocytosis of nidogens as well as H_C_T. Indeed, addition of recombinant FNIII domains at concentrations equivalent to one-tenth of their *in vitro* binding affinities (0.25 μM), was nearly equally efficient at reducing nidogen and H_C_T uptake as were concentrations ten times higher their binding affinities (20 μM). In contrast, addition of a single recombinant FNIII domain had limited effect on H_C_T and nidogen internalisation. Whilst addition of the 5^th^ FNIII domain did not result in any change, the 2^nd^, 4^th^ and 7^th^ FNIII domains yielded a modest reduction in nidogen-H_C_T complex internalisation, underscoring that the LAR-nidogen interaction is driven by the overall avidity of FNIII domains to nidogens.

Interestingly, the decrease in endocytosed nidogen-H_C_T levels seen with the addition of multiple FNIII domains (~45-50%) corresponds well with that observed upon LAR knockdown using shRNA (~40-45%). The incomplete blocking of H_C_T internalisation under these experimental conditions indicates that the nidogen-TeNT complex relies on additional membrane proteins for its endocytosis. Our study suggests that PTPRδ is a strong candidate for this function, given that it binds to both nidogen-1 and −2 and shares significant sequence similarity with LAR. Concomitantly, it was also found to be a key player in the TrkB signalling cascade in cortical neurons (Tomita *et al*, 2020).

Taken together, our results provide the first identification of a membrane receptor complex for TeNT at the NMJ. By identifying LAR as a binding partner of nidogens, which enables their internalisation, we have revealed a pathway by which TeNT is targeted to axonal signalling endosomes and undergoes longdistance transport to the neuronal cell body. The LAR-nidogen complex, along with polysialogangliosides, thus acts as an efficient capture mechanism for TeNT at the NMJ, which enables its binding at very low concentrations and ferries it into motor neurons. Whilst ECM proteins have been reported to bind to a multitude of surface molecules, this is the first report of a membrane-bound receptor enabling the internalisation and trafficking of an ECM protein in neurons, the physiological relevance of which remains to be uncovered. We have also established a molecular link between the ECM and trophic pathways in the nervous system, suggesting that nidogens might play critical roles in controlling growth factor availability at synapses as well as regulating neuronal signalling.

## Materials and methods

### Key resources

**Table 1.** Reagents.

| Name | Source | Identifier |
| --- | --- | --- |
| Anti-HA magnetic beads | Thermo Fisher Scientific | 88837 |
| AlexaFluor 555 C2 maleimide | Thermo Fisher Scientific | A20346 |
| AlexaFluor 647 C2 maleimide | Thermo Fisher Scientific | A20347 |
| Brain-derived neurotrophic factor (BDNF) | Peprtech | 450-02-100 |
| Ciliary neurotrophic factor (CNTF) | Peprtech | 450-50-50 |
| Coomassie brilliant blue R-250 | Bio-Rad Laboratories | 1610400 |
| Dulbecco's modified Eagle medium (DMEM) | Gibco | 41966-029 |
| Foetal bovine serum (FBS) | Thermo Fisher Scientific | 10309433 |
| Gibson assembly cloning kit | New England Biolabs | E5510S |
| Glial-derived neurotrophic factor (GDNF) | Peprtech | 450-10-50 |
| GlutaMAX | Thermo Fisher Scientific | 35050061 |
| iMatrix-511 | Reprocell | NP892-011 |
| Laminin | Merck | L2020 |
| Lenti-X concentrator | Takara Bio | 631232 |
| Lipofectamine 3000 | Invitrogen | L3000008 |
| Neurobasal medium | Thermo Fisher Scientific | 21103049 |
| NeuroMag transfection reagent | OZ Biosciences | NM50200 |
| Opti-MEM reduced media | Thermo Fisher Scientific | 31985062 |
| Penicillin-streptomycin | Thermo Fisher Scientific | 15-140-122 |
| PF-06273340 | Merck | PZ0254 |
| Poly-L-ornithine | Merck | P4957 |
| Protein A sepharose(R) 4B, fast flow | Merck | P9424 |
| Protein G sepharose 4, fast flow | Merck | GE17-0618-01 |
| Recombinant mouse nidogen-2 protein | R&D Systems | 6760-ND-050 |
| Retinoic acid | Merck | R2625-50 |
| 1-Step ultra TMB solution | Thermo Fisher Scientific | 34029 |

**Table 2.**
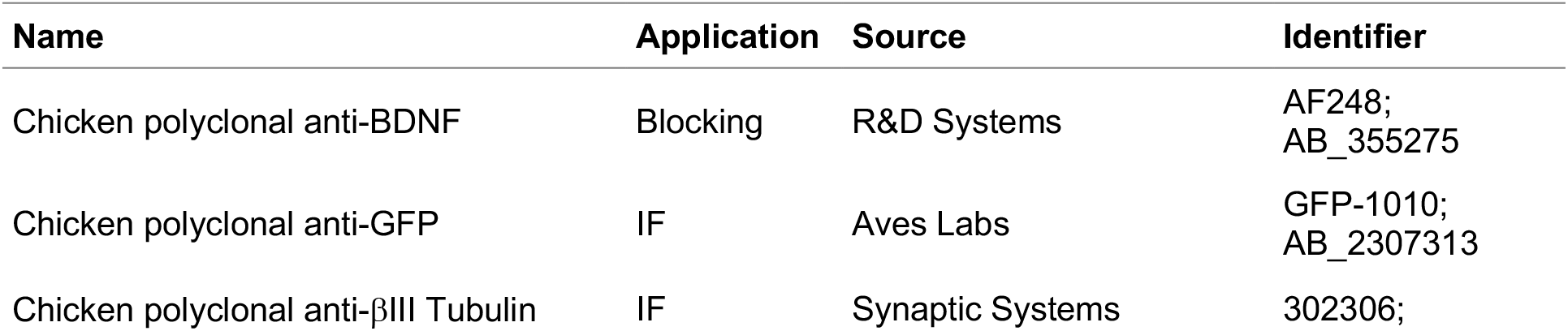
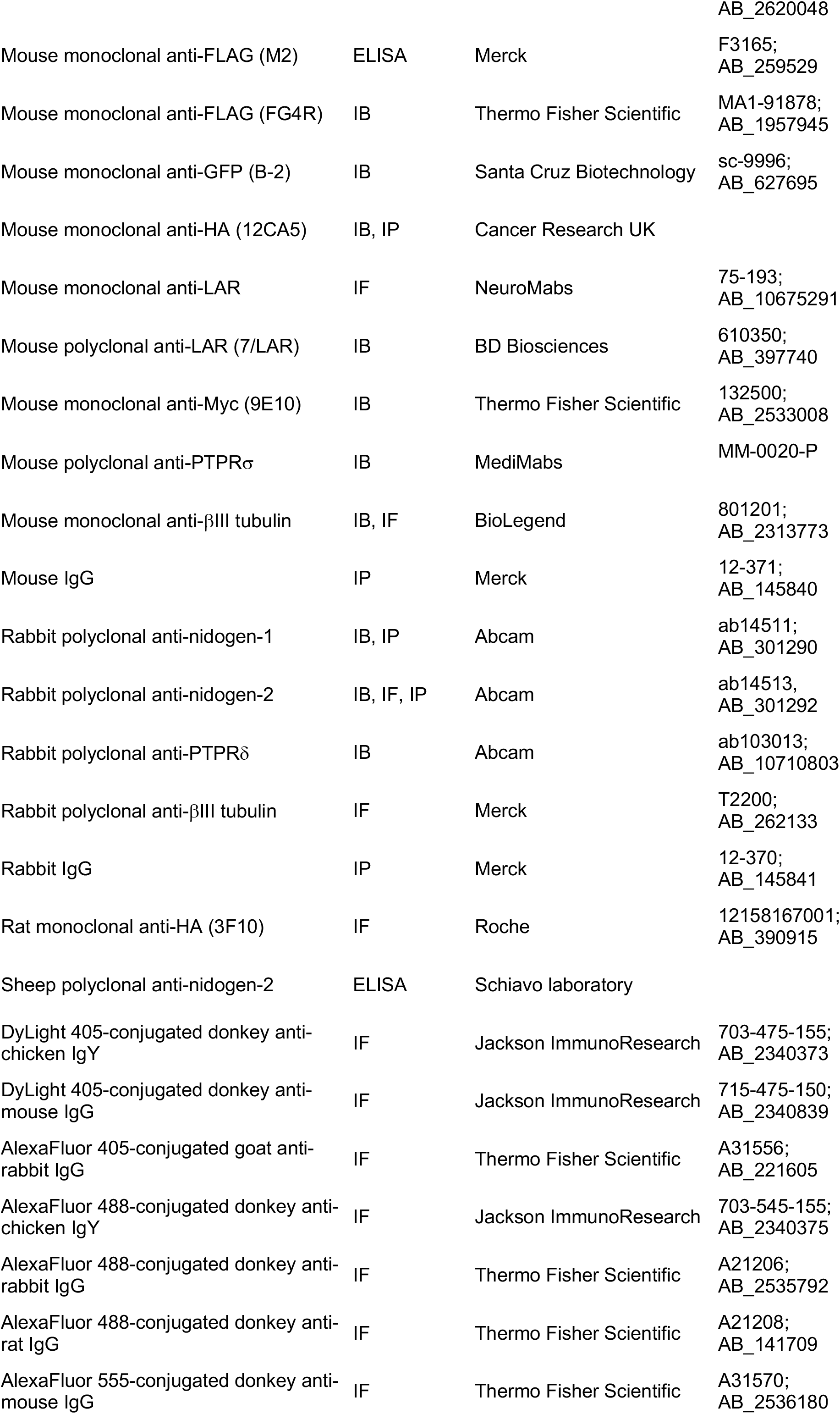
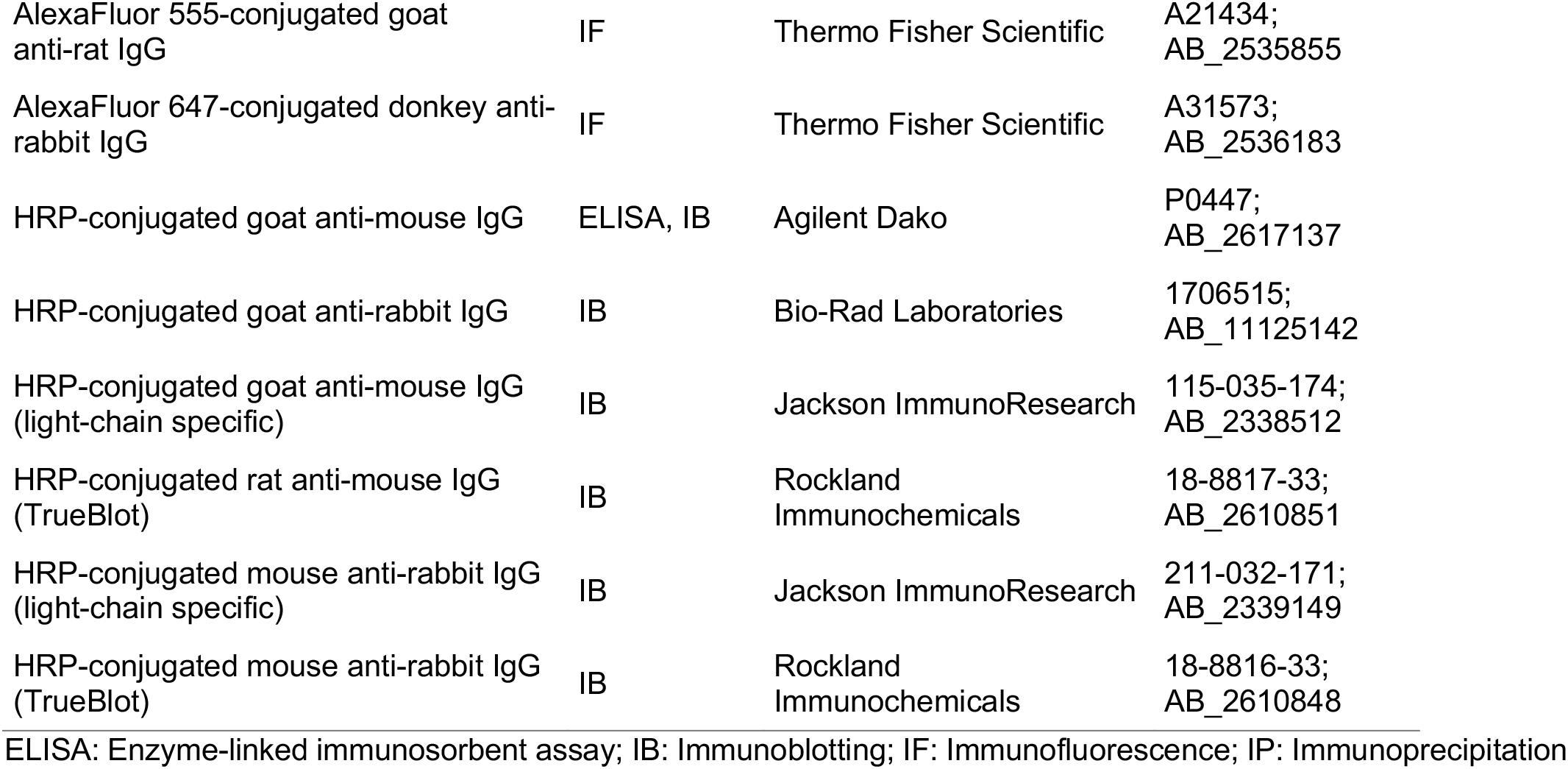
Primary and secondary antibodies.

### Plasmids and cloning

pDisplay plasmids encoding HA-eLAR-Myc, HA-ePTPRδ-Myc and HA-ePTPRσ-Myc were kindly provided by the Sala laboratory (University of Milan, Italy) (Valnegri *et al*, 2011). In these constructs, the extracellular domain of each human RPTP is fused to the murine Igκ chain leader sequence and an HA tag (YPYDVPDYA) at the N-terminus. The C-terminus is fused to a Myc tag (EQKLISEEDL) and the platelet-derived growth factor (PDGF) receptor transmembrane domain, enabling surface localisation of the expressed proteins. pCEP.Pu plasmids encoding mouse nidogen-1 and −2 were provided by Dr. Takako Sasaki (Oita University, Japan) (Bechtel *et al*, 2012). For this study, nidogen-2 was cloned into a pcDNA3.1 vector and tagged with a 6×His tag at the C-terminus. pDisplay plasmids containing individual LAR domains were made using the pDisplay-HA-eLAR-Myc plasmid. For bacterial expression and purification, the 2^nd^, 4^th^, 5^th^ and 7^th^ FNIII domains of LAR were cloned into the pET28a(+) vector, containing an N-terminal 6×His tag and a C-terminal FLAG tag (DYKDDDDK). Cloning was done using standard Gibson or inverse PCR cloning strategies and positive clones were selected using colony PCR and Sanger sequencing. Plasmids encoding LAR shRNA and scrambled controls were purchased from GeneCopoeia. Packaging (pCMVR8.74; Addgene plasmid no. 22036; RRID: Addgene_22036) and envelope plasmids (pMD2.G; Addgene plasmid no. 12259; RRID: Addgene_12259) were originally prepared by the Didier Trono laboratory (École Polytechnique Fédérale de Lausanne, Switzerland).

### Cell lines

N2a cells were sourced from Cancer Research UK London Research Institute Cell Services, whilst Lenti-X HEK293T cells were sourced from ClonTech. Both cell lines were cultured in DMEM with 10% FBS and 1% GlutaMAX. Cells were split every 2–3 days at 80-90% confluency. For immunoprecipitation experiments, N2a cells were plated on Petri dishes, cultured for 1 day *in vitro* (DIV), and then differentiated using 10 μM retinoic acid for 48-72 h. For immunofluorescence experiments, they were plated on poly-D-lysine coated coverslips and cultured for 2 DIV. Lenti-X HEK293T were plated directly on Petri dishes for lentiviral production.

### Motor neuron cultures

Mixed embryonic ventral horn cultures, referred to in this study as primary motor neurons, were isolated from E11.5-13.5 mouse embryos as previously described (Fellows *et al*, 2020). Briefly, ventral horns from E11.5-13.5 pregnant wild-type mice (C57/Bl6, Charles River) were dissociated, centrifuged at 380*g* for 5 min, seeded on poly-L-ornithine- and laminin-coated coverslips or wells, and maintained in motor neuron media (Neurobasal with 2% v/v B27, 2% heat-inactivated horse serum, 1x GlutaMAX, 24.8 μM β-mercaptoethanol, 10 ng/ml CNTF, 0.1 ng/ml GDNF, 1 ng/ml BDNF, 1x penicillin-streptomycin) at 37°C and 5% CO_2_.

### Lentiviral particle production and motor neuron transduction

LAR shRNA and control viral particles were generated by co-transfecting shRNA, packaging and envelope plasmids into Lenti-X HEK293T cells with Lipofectamine 3000 using manufacturer’s instructions. Medium containing lentiviral particles was collected at 48 and 72 h after transfection, concentrated using Lenti-X concentrator and resuspended in Opti-MEM media. shRNA viral particles were stored at −80 °C until needed. Neurons were transduced on DIV3 by adding viral particles directly to the medium. After 48 h (DIV5), motor neurons were either lysed for western blot analyses or immunostained.

### H_C_T preparation and labelling

HA-H_C_T, VSVG-H_C_T and H_C_T fused to a cysteine-rich tag were prepared as previously described (Restani *et al*, 2012). HA-H_C_T was used in immunofluorescence, whilst VSVG-H_C_T in immunoprecipitations. H_C_T fused to a cysteine-rich tag was labelled with AlexaFluor 555 C2 maleimide or AlexaFluor 647 C2 maleimide following manufacturer’s instructions and used in direct immunofluorescence assays.

### Cell-based assays and immunofluorescence

General immunofluorescence protocols were carried out as described below. Ventral horn cultures at DIV5, after appropriate treatment and incubation, were cooled on ice. Surface-bound probes were removed by washing the cells in mildly acidic buffer (0.2 M acetic acid, 0.5 M NaCl, pH 2.4) for 1 min on ice. After a wash with cold PBS, cells were fixed (4% paraformaldehyde and 5% sucrose in PBS), blocked and permeabilised (10% horse serum, 0.5% bovine serum albumin and 0.2% Triton X-100 in PBS) for 10 min at room temperature. Cells were then stained with fluorescently labelled primary and secondary antibodies in blocking buffer for 1 h each at room temperature and mounted. All primary and secondary antibodies were used at 1:500 and 1:1,000, respectively.

For surface expression/localisation analyses, N2a cells were transfected with the appropriate plasmids at DIV1. After 24 h, cells were washed with PBS and treated with an anti-HA or anti-nidogen antibody in blocking buffer (without Triton X-100) for 1 h at room temperature. After another wash with PBS, cells were fixed, permeabilised and taken forward for immunostaining using AlexaFluor 488-conjugated anti-rat or anti-rabbit antibodies. Nuclei were stained by supplementing the secondary antibody solution with 0.5 μg/ml 4’,6-diamidino-2-phenylindole.

For co-localisation and correlation experiments, motor neurons plated on poly-L-ornithine and iMatrix 511-coated coverslips were incubated with 40 nM H_C_T-647 and an anti-nidogen-2 antibody (1:500) for 1 h at 37°C. After acid washing, fixation and permeabilization, cells were stained with primary antibodies against LAR and βIII tubulin, which were revealed using an AlexaFluor 555-conjugated anti-mouse antibody and DyLight 405-conjugated anti-chicken antibody, respectively. Internalised nidogen-2 was revealed using an AlexaFluor 488-conjugated anti-rabbit antibody.

For LAR knockdown, ventral horn cultures were transduced with lentiviral particles on DIV3. Motor neurons transduced with LAR shRNA#2-expressing lentiviruses, were magnetofected with 0.5 μg of pDisplay-HA-eLAR-Myc plasmid on DIV4 to express the HA-eLAR-Myc fusion protein, thus rescuing endogenous LAR depletion. On DIV5, cultures were incubated with 25 nM H_C_T-647 for 1 h at 37°C, acid washed and then stained as described above. Anti-βIII tubulin, anti-GFP and anti-HA antibodies were used for immunodetection, which were revealed using DyLight 405-conjugated anti-rabbit, AlexaFluor 488-conjugated anti-chicken and AlexaFluor 555-conjugated anti-rat secondary antibodies, respectively.

To check the relevance of TrkB activity on H_C_T internalisation, motor neuron media was replaced by Neurobasal for 1 h. Motor neuron media was supplemented with 100 nM PF-06273340 and an anti-BDNF blocking antibody (1:50) before being added back to the cells. After 30 min, HA-H_C_T (25 nM) was added, incubated for 1 h at 37°C, followed by immunostaining using anti-HA and βIII tubulin antibodies. Primary antibodies were revealed using AlexaFluor 488-conjugated anti-rat and AlexaFluor 647-conjugated antirabbit antibodies, respectively.

For H_C_T uptake experiments in the presence of recombinant LAR domains, the individual 2^nd^, 4^th^, 5^th^ and 7^th^ LAR FNIII domains were cloned, expressed and purified (Vilstrup *et al*, 2020). Purified proteins were dialyzed (25 mM Tris-Cl pH 7.4, 125 mM NaCl, 5% glycerol) and stored at −20°C. Motor neuron cultures were pulsed with 40 nM H_C_T-555 and a nidogen-2 antibody (1:500), in the presence of recombinant LAR fibronectin III domains for 30 min on ice. After replacement of media with fresh motor neuron media, cultures were shifted to 37°C and allowed to internalise the nidogen-H_C_T complex for 45 min. This was followed by acid washing and immunodetection using a mouse anti-βIII tubulin antibody. DyLight 405-conjugated anti-mouse and AlexaFluor 488-conjugated anti-rabbit secondary antibodies were used to reveal the cellular localisation of βIII tubulin and internalised nidogen-2, respectively.

### Image acquisition and analysis

All images were acquired using an inverted Zeiss LSM 780 (with a 40x, 1.3 NA DIC Plan-Apochromat oil-immersion objective) or Zeiss LSM 980 (with a 40x, 1.3 NA DIC Plan-Neofluar oil-immersion objective) confocal microscope. Co-localisation experiments were imaged on Zeiss LSM 980 in airy-scan mode, using a 63x, 1.4 NA DIC Plan-Apochromat oil-immersion objective. All related images were acquired under the same microscope settings, processed using Fiji, and, when appropriate, scaled using the same settings. Maximum intensity-projected z stacks were used for image analysis, which was performed in SynPAnal (Danielson & Lee, 2014). βIII tubulin-positive neurites were manually selected, thresholded, and integrated intensities per unit of axonal length of internalised H_C_T and nidogen-2 were quantified. In LAR knockdown experiments, GFP was used as a marker of lentiviral transduction, hence neurites positive for both GFP and βIII tubulin were selected. In rescue experiments with recombinant HA-eLAR-Myc, neurites positive for HA, GFP and βIII tubulin were selected for analysis. Co-localisation and correlation analysis were performed in Fiji using in-built Plot Profile and intensity measurement functions, respectively. In correlation analysis, 3-4 neurites of 35-60 μm length were randomly selected in each image in the βIII tubulin channel and corresponding integrated intensities of H_C_T-647, LAR and nidogen-2 were calculated.

### Cell lysis and immunoprecipitations

Relevant plasmids were transfected in N2a cells (DIV1) using Lipofectamine 3000 and manufacturer’s protocols. After 6 h, culture media was replaced with differentiation media (DMEM, 1% FBS, 1% GlutaMAX, 10 μM retinoic acid). After 48-72 h, cells were treated with 80 nM VSVG-H_C_T for 10 min at 37°C, and then lysed in immunoprecipitation buffer (20 mM Tris-Cl pH 8.0, 137 mM NaCl, 10% glycerol, 0.5% NP-40, protease and phosphatase inhibitors) for 20 min on ice. Cell debris was removed by centrifuging the lysates at 21,000*g* for 15 min at 4°C. Two types of beads were used for immunoprecipitation experiments: i) Protein A or G sepharose beads non-covalently bound to appropriate antibodies for 2 h at 4°C (**Figure 1**, **Figure 1 - figure supplement 1**); or, ii) Magnetic beads covalently conjugated to an anti-HA antibody (**Figure 4**). Lysates were applied to beads for 2 h at 4°C under gentle agitation, washed 3-5 times in immunoprecipitation buffer and then treated in 4× Laemmli buffer (250 mM Tris-Cl pH 6.8, 8% SDS, 40% glycerol, 0.02% bromophenol blue, 10% β-mercaptoethanol) at 95°C for 4 min. Samples were analysed by western blotting. Relevant controls were used for each experiment, as mentioned in figure legends. 5% lysate was loaded as input.

Motor neuron lysates were prepared in RIPA buffer (50 mM Tris-Cl pH 7.5, 150 mM NaCl, 1% NP-40, 0.5% sodium deoxycholate, 0.1% SDS, 1 mM EDTA, 1 mM EGTA, protease and phosphatase inhibitors) as described above and analysed using western blotting. Total protein was probed using Coomassie R-250 staining on membranes. All blots were imaged on the ChemiDoc™ Touch Imaging System (Bio-Rad Laboratories), whilst data analysis was performed using the ImageLab software (Bio-Rad Laboratories).

### Enzyme-linked immunosorbent assays

1 mg/ml anti-nidogen-2 capture antibody, diluted in PBS, was applied to wells, gently agitated for 5 minutes at room temperature, and incubated overnight at 4°C. Wells were blocked using 5% bovine serum albumin in PBS containing 0.05% Tween-20 (PBST) for 1.5 h at room temperature. 0.5 picomoles of recombinant nidogen-2 was mixed with varying concentrations of bacterially expressed LAR FNIII domains (50 nM-30 μM) in binding buffer (20 mM Tris-Cl pH 8.0, 137 mM NaCl, 10% glycerol, 0.5% NP-40) and shaken for 2 h at 4°C. The mixture was applied to the wells and allowed to bind for 1.5 h at room temperature. After multiple washes using PBST, the bound LAR FNIIIx-nidogen-2 complex was detected using an anti-FLAG primary antibody (1:1,000) and an HRP-conjugated anti-mouse secondary antibody (1:5,000). Both antibodies were diluted in PBST containing 1% bovine serum albumin and incubated for 1 h at room temperature. 3,3’,5,5’-tetramethylbenzidine was used as a substrate and the reaction was stopped using 2 M H_2_SO_4_. Complex formation was assessed by measuring absorbance at 450 nm on a multi-mode FLUOstar Omega microplate reader (BMG Labtech). Reactions devoid of any LAR FNIII domain served as background. All readings in an experiment were normalised to absorbance values obtained with 5 μM LAR FNIII.

### Statistical analyses

Data were tested for normality using the Kolmogorov-Smirnov test, while equal variance between groups was assumed. Normally-distributed data were statistically analysed using unpaired *t*-test or one-way analysis of variance (ANOVA) followed by Dunnett’s multiple comparisons test. Non-normally distributed data were analysed using Kruskal-Wallis test followed by Dunn’s *post-hoc* test. Means ± standard error of the mean were plotted. All tests were two-sided and an α-level of *P* < 0.05 was used to determine significance. All *P* values are indicated in the figures and/or figure legends. GraphPad Prism 9 software (version 9.5.0) was used for statistical analyses and figure production.

## Ethics statement

All experiments were conducted under the guidelines of the UCL Queen Square Institute of Neurology Genetic Manipulation and Ethics Committees and in accordance with the European Community Council Directive of 24 November 1986 (86/609/EEC). Animal experiments were performed under license from the UK Home Office in accordance with the Animals (Scientific Procedures) Act, 1986 and were approved by the UCL Queen Square Institute of Neurology Ethical Review Committee.

## Acknowledgements

We thank Prof. Carlo Sala (University of Milan, Italy) for LAR, PTPRδ and PTPRσ plasmids, the personnel of the Denny Brown Laboratories (Queen Square Institute of Neurology, University College London, UK) for maintenance of mouse colonies, Dr. Darius Vasco Köster (Warwick University, UK) for thoughtful discussions, and Drs. James N. Sleigh and Elena R. Rhymes (Queen Square Institute of Neurology, University College London, UK) for critical reading of the manuscript. We thank Simon L. Duerr (École Polytechnique Fédérale de Lausanne, Switzerland) for generously sharing high-quality science illustrations *via* https://bioicons.com/, which were used for rendering schematics in this manuscript. This project was funded by a Human Frontier Science Program Long-Term Fellowship LT000220/2017-L (SS), Wellcome Trust Awards 107116/Z/15/Z and 223022/Z/21/Z (GS), UK Dementia Research Institute Foundation Award UKDRI-1005 (GS), and an Alzheimer’s Society PhD Studentship Grant 520 (GS).

## Author contributions

SS and GS conceived the work and designed the experiments; SS, DV-C, CP, SSN and SR performed the experiments; GZ performed structural analyses of the nidogen-LAR complex; SS and GS analysed the data; SS produced the figures and wrote the paper, with input from all authors; SS and GS acquired funding for this study. All authors have approved submission of this work. The funders had no role in study design, data collection and analysis, decision to publish, or manuscript preparation.

## Conflict of interest

The authors have no conflicts of interest to declare.

## Supplementary Information

**Supplementary Figure 1.**
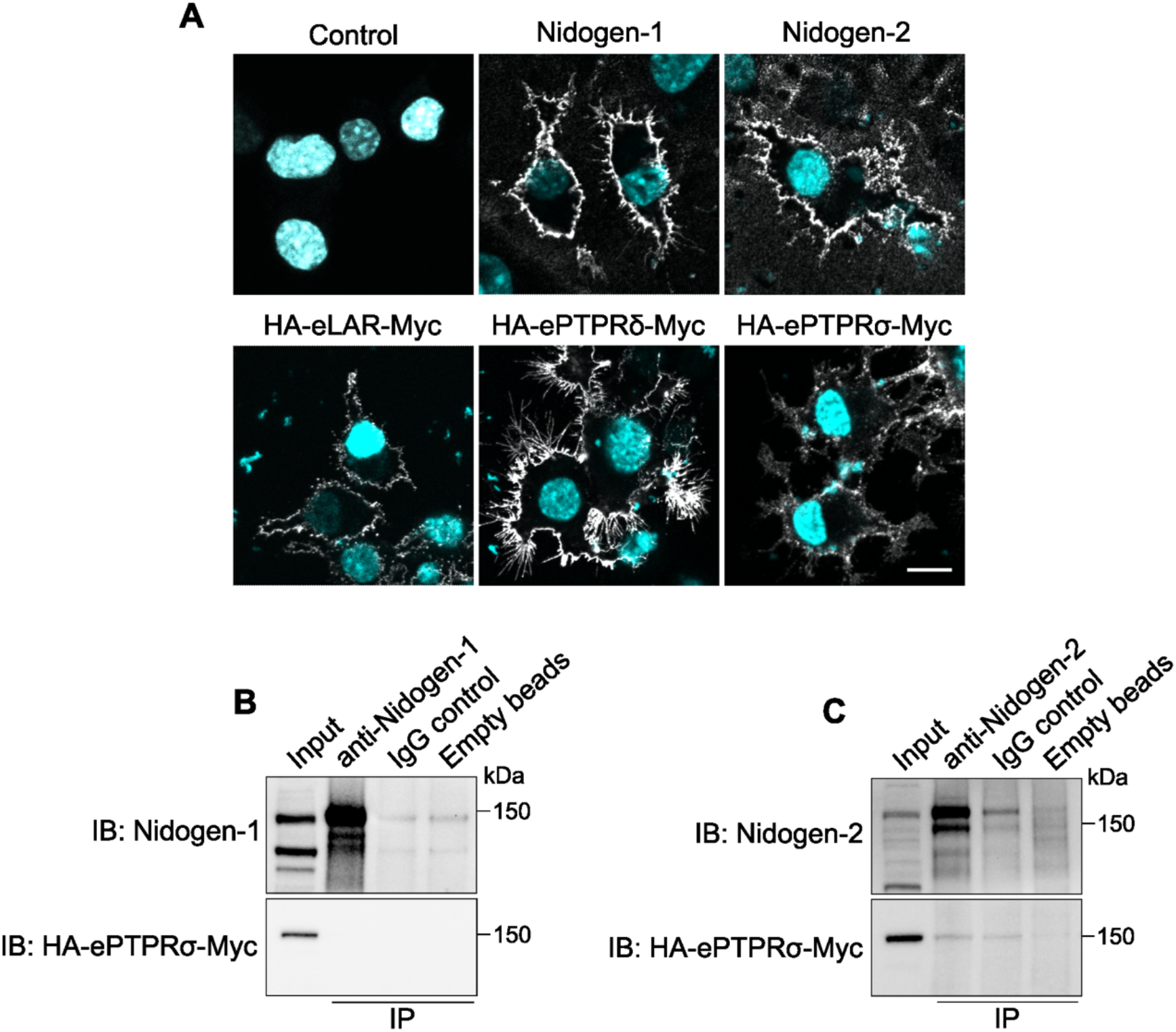
**(A)** Representative immunofluorescence images showing localisation of recombinant nidogen-1, nidogen-2, eLAR, ePTPRδ and ePTPRσ in mouse N2a cells upon transient expression. Nuclei stained with 4’,6-diamidino-2-phenylindole have been pseudo-coloured in cyan. Scale bar: 5 μm. **(B, C)** Unlike LAR and PTPRδ, PTPRσ does not interact with nidogens in the presence of H_C_T. Nidogens were immunoprecipitated from N2a cell lysates, which were then probed for HA-ePTPRσ-Myc using an anti-HA antibody. Non-specific antibodies bound to beads and empty beads were used as negative controls; 5% input was loaded.

**Supplementary Figure 2.**
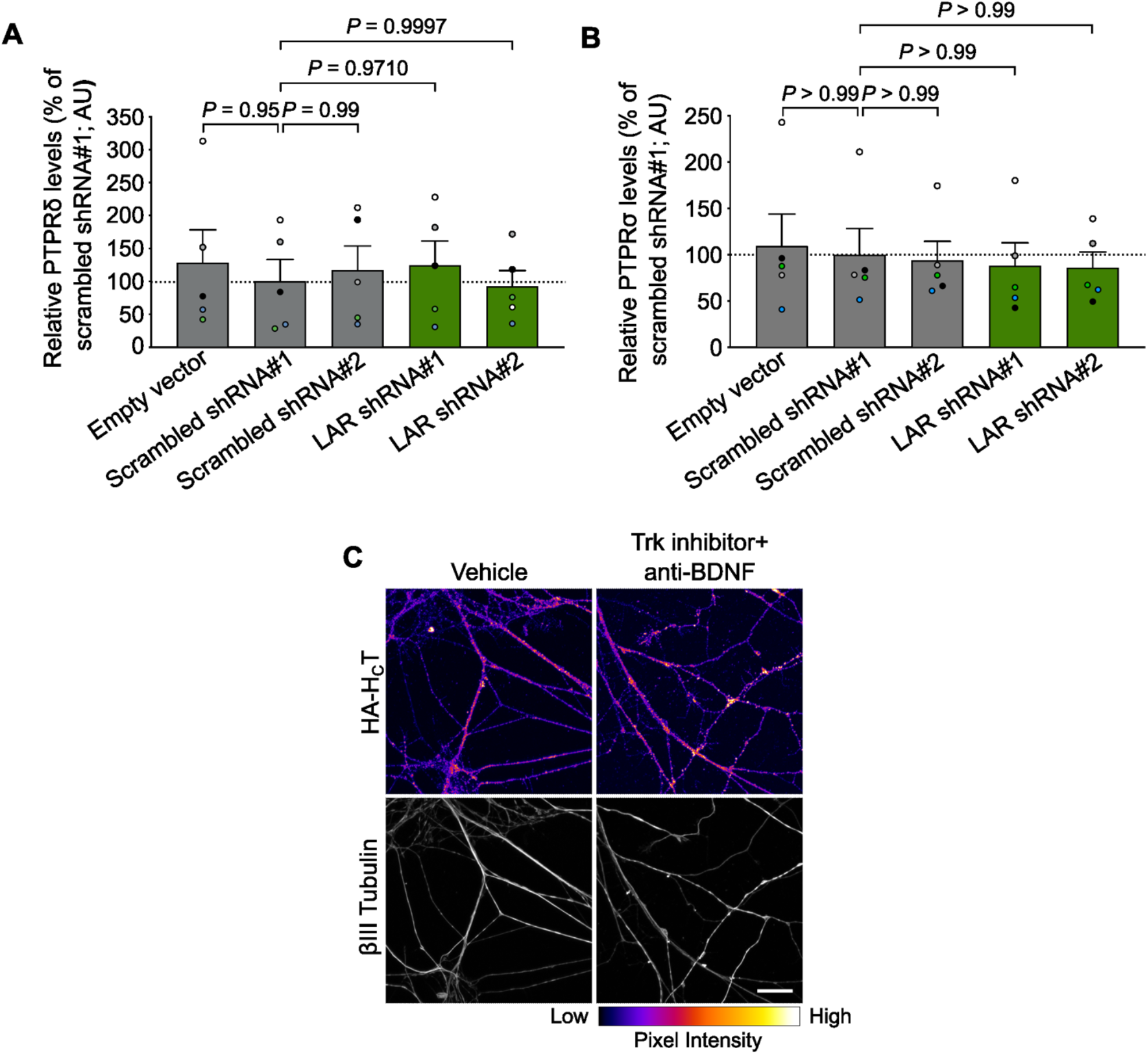
**(A)** PTPRδ quantification in lysates of ventral horn cultures transduced with lentiviruses encoding shRNAs against mouse LAR. Results were tested for statistical significance using one-way ANOVA (*P* = 0.9480), followed by Dunnett’s multiple comparisons test (n = 5 independent experiments; error bars indicate s.e.m.). **(B)** PTPRσ quantification in lysates of ventral horn cultures transduced with lentiviruses encoding shRNAs against mouse LAR. Results were tested for statistical significance using Kruskal-Wallis test (*P* = 0.9665), followed by Dunn’s *post-hoc* test (n = 5 independent experiments; error bars indicate s.e.m.). Data are presented as a percentage of the levels of PTPRδ (A) and PTPRσ (B) in neurons treated with scrambled shRNA#1. Lentiviruses carrying an empty vector and two scrambled shRNAs were used as negative controls. **(C)** Representative confocal images of endocytosed HA-H_C_T in motor neurons treated with the pan-Trk inhibitor PF-06273340 and an anti-BDNF antibody. Images in the top panel have been colour mapped based on their intensities. Vehicle refers to DMSO-treated cultures. Scale bar: 20 μm.

**Supplementary Figure 3.**
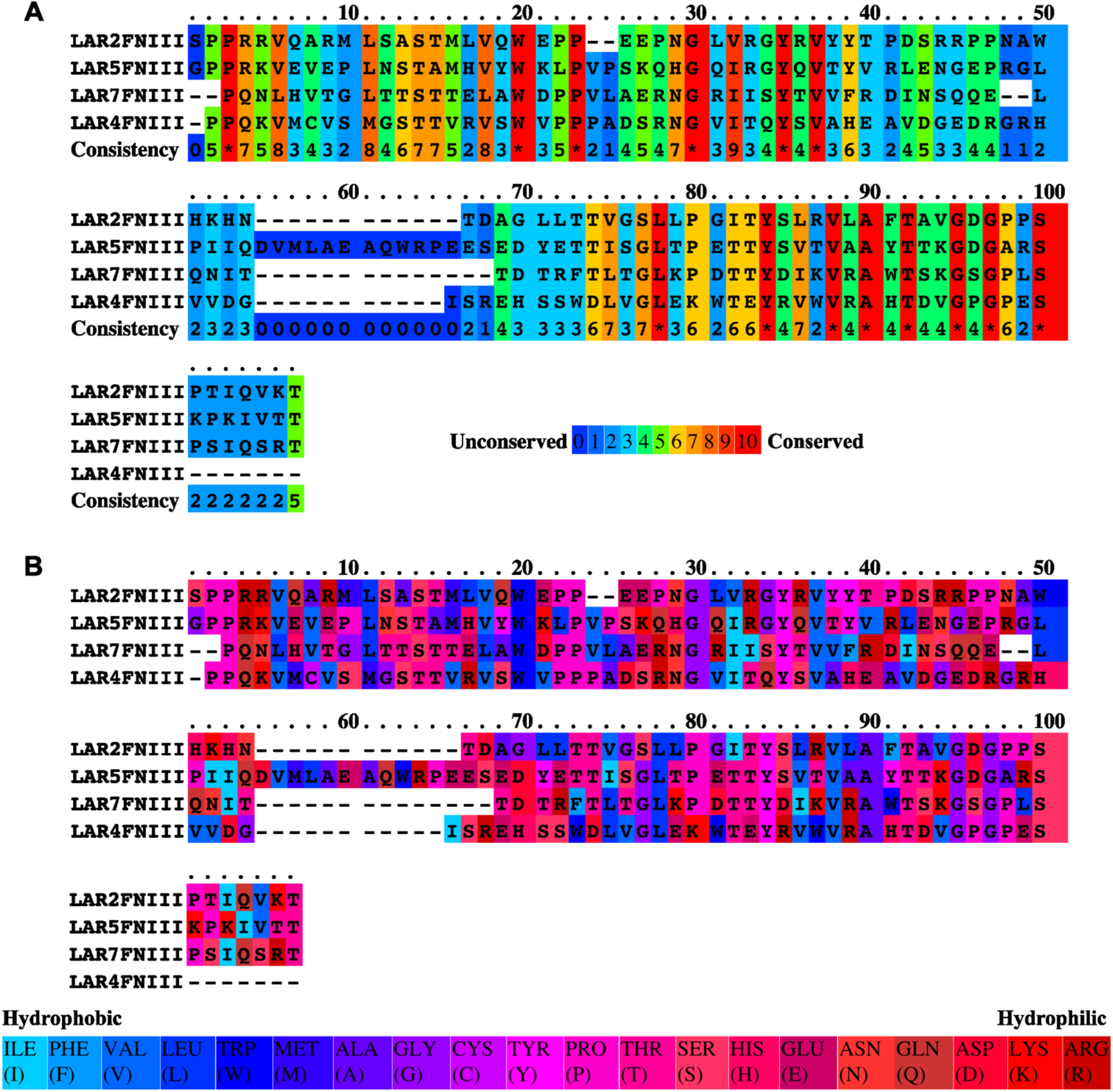
**(A)** Sequence alignment of the human LAR FNIII2, FNIII4, FNIII5 and FNIII7 domains. Scoring ranges from 0 for the least conserved alignment position, up to 10 for the most conserved position. **(B)** Assessment of conservation of hydrophobicity/hydrophilicity in nidogen-binding domains of LAR. Colour assignments from hydrophobic to hydrophilic are presented below the alignment.

**Supplementary Figure 4.**
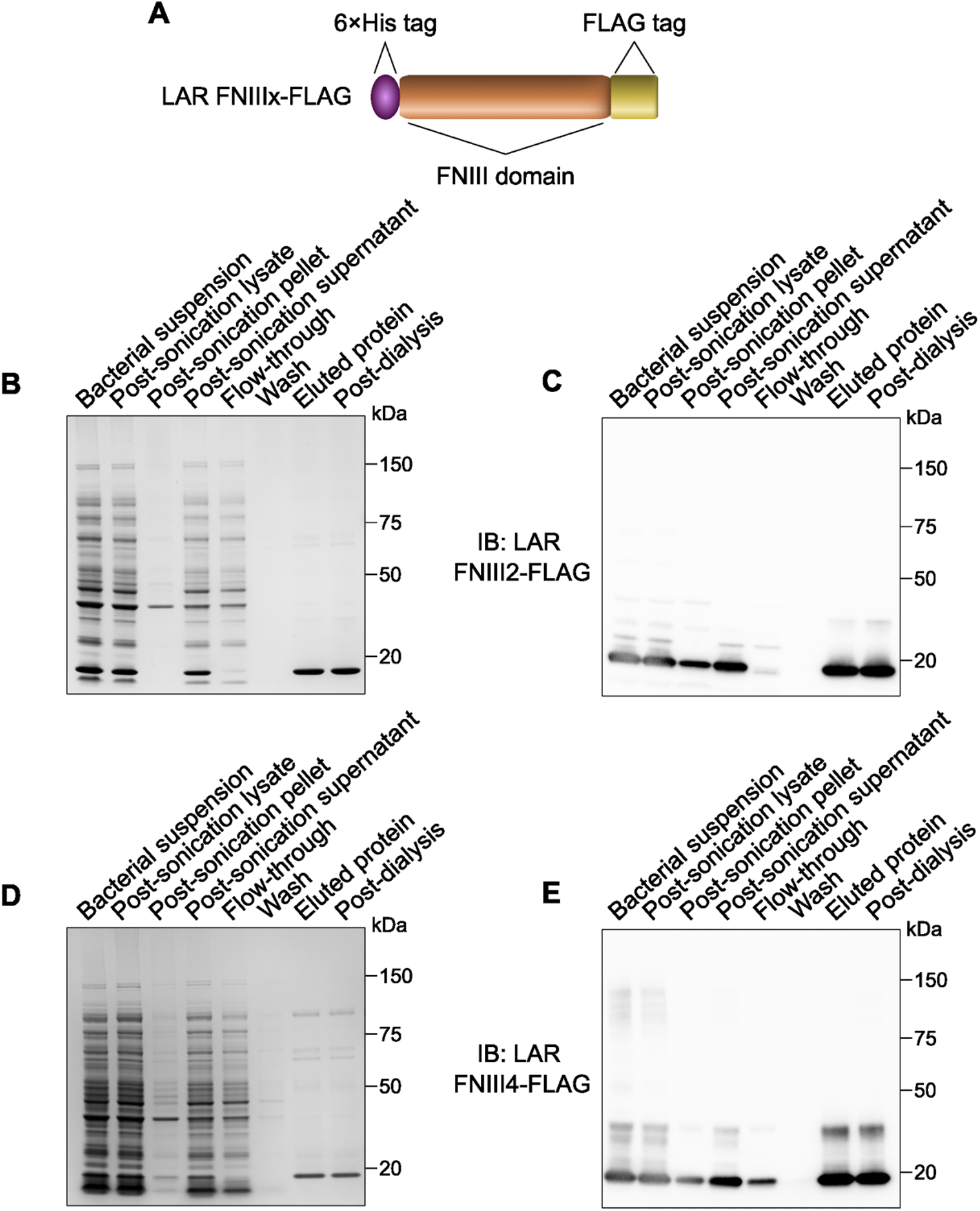
**(A)** Schematic of LAR FIII fusion proteins used for bacterial expression and purification. Each nidogen-binding FNIII domain was tagged with a 6×His tag at the N-terminus and a FLAG tag at the C-terminus. **(B, C)** Coomassie staining (B) and western blot (C) showing bacterial expression and Ni^2+^-affinity purification of LAR FNIII2-FLAG (predicted molecular weight is ~15 kDa). **(D, E)** Coomassie staining (D) and western blot (E) showing bacterial expression and Ni^2+^-affinity purification of LAR FNIII4-FLAG (predicted molecular weight is ~15 kDa). Western blots were probed using an anti-FLAG antibody.

**Supplementary Figure 5.**
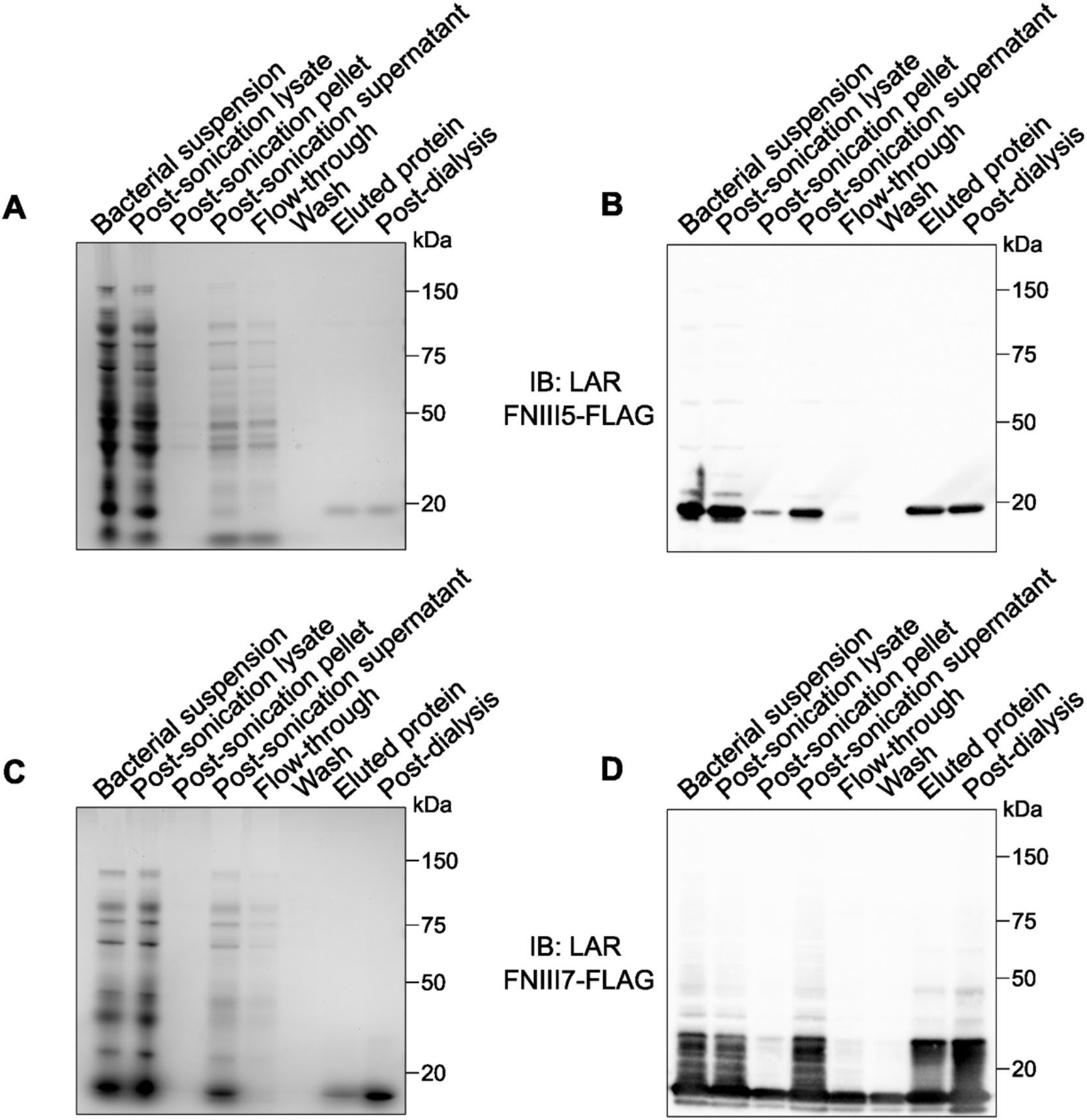
**(A, B)** Coomassie staining (A) and western blot (B) showing bacterial expression and Nibaffinity purification of LAR FNIII5-FLAG (predicted molecular weight is ~17 kDa). **(C, D)** Coomassie staining (C) and western blot (D) showing bacterial expression and Ni^2+^-affinity purification of LAR FNIII7-FLAG (predicted molecular weight is ~15 kDa). Western blots were probed using an anti-FLAG antibody.

## Notes

### Competing Interest Statement

The authors have declared no competing interest.

